# Hippocampal circuit-specific enhancement of GABA-inhibition caused by discrete gene regions in a Down syndrome model

**DOI:** 10.1101/2025.07.21.665861

**Authors:** Saad B Hannan, Eva Lana-Elola, Sheona Watson-Scales, Elizabeth M.C. Fisher, Victor L.J. Tybulewicz, Trevor G. Smart

## Abstract

Although Down syndrome (DS), trisomy 21, affects ∼6 million people worldwide, the neural circuit mechanisms underlying the neurophenotypes of impaired learning, memory and language are unknown. A prominent candidate mechanism involves dysfunctional GABA-signalling and GABA_A_ receptor ligands have been proposed as therapeutics to reverse the neurophenotypic effects of DS.

By investigating GABA neurotransmission in brain regions important for cognition in mouse DS models, we reveal that excessive inhibition is not a ubiquitous feature of DS but instead is brain circuit-specific demonstrating increased phasic and tonic inhibition in the dentate gyrus with no comparative changes to inhibition in CA1 and medial prefrontal cortex.

In the dentate, elevated extrasynaptic GABA signalling, and interneuron numbers, likely underpin spike firing defects. We show that increased GABA inhibition is caused by increased dosage of *Olig1*, *Olig2* and *Dyrk1a*. Overall, DS mice are characterised by circuit-specific dysfunctional inhibition predicted to affect cognition via sparse coding in the hippocampus.

## Introduction

Down syndrome (DS), is the most common form of genetically-driven intellectual disability affecting 1 in 800 births^1^. It is caused by trisomy of the smallest human chromosome, 21 (Hsa21). DS results in multiple phenotypes, driven by an extra copy of one or more of the 221 coding genes and other genetic elements on Hsa21. One of the most prevalent DS phenotypes is cognitive impairment, which affects all DS individuals. A long-standing hypothesis is that the cognitive deficits are caused by increased neuronal signalling by the neurotransmitter γ-aminobutyric acid (GABA)^2–4^ acting via GABA_A_ receptors which are pseudo-symmetric heteropentamers^5–8^ formed from α1-6, β1-3, γ1-3, δ, ε, π, θ and ρ1-3 subunits to yield receptors that are dominated by generic αβγ and αβδ isoforms displaying diverse functional and pharmacological properties^9^.

Physiologically, these specific GABA_A_R subtypes operate over different timescales, as a consequence of varying sources of GABA^10,11^, to impart phasic (synaptic) and tonic (ambient) inhibition of neural excitability at discrete locations of the postsynaptic neuronal membrane^12^. Interestingly, brain-area specific changes to tonic inhibition, in particular, underlie multiple neurological disorders such as absence epilepsy, Alzheimer’s disease, Rett syndrome and anxiety^13–16^.

The importance of GABA-signalling for cognitive deficiency in DS originates from observations that inhibition is markedly increased in Ts65Dn mice, an early mouse model of DS^2,17,18^. This strain has an extra copy of 132 Hsa21-orthologous genes, thereby partially modelling Hsa21 trisomy^19^. In this model, reversal of elevated signalling by an antagonist or a negative allosteric modulator at GABA_A_ receptors recalibrated impaired cognition and synaptic plasticity in the hippocampus, which is a cellular correlate of learning and memory^17–22^. Surprisingly, the cognitive benefit of reducing GABA signalling in Ts65Dn considerably exceeded the bioavailability of the GABA antagonist suggesting far-reaching benefits presumably involving long-term neural circuit plasticity and/or rewiring.

Since these initial observations, GABA signalling has been further implicated in DS cognitive impairment evidenced by increased interneuron and/ or inhibitory synapse numbers in Ts65Dn forebrain^23,24^ and hippocampus^19,25–27^ with an increase in the rate and magnitude of GABA-mediated synaptic input into Ts65Dn hippocampus^24,29,30^. Similarly, increased interneuron numbers have been reported in human organoids derived from DS donors^28^. These cellular and synaptic changes may underlie perturbations to neural circuitry, neurophysiology and synaptic plasticity, since long-term potentiation (LTP) is reduced^17,18,26,29–38^ and long-term depression (LTD) is elevated^39^ in multiple mouse models. Moreover, application of GABA_A_ receptor antagonists correct LTP deficits in Ts1Cje^38^ and Ts1Rhr^36^ mouse DS models.

Unexpectedly, in adult CA1 Ts65Dn neurons, GABA has been reported to be excitatory^40^ although a previous study^41^ reported GABA to be inhibitory in the same cells at earlier ages, when GABA is more likely to be depolarising. Nevertheless, taking account of studies so far, there is a broad consensus that over-active GABA transmission is a unifying mechanism underlying cognitive deficits in DS. Thus it was surprising when a recent clinical trial using a GABA_A_ receptor α5 receptor-specific negative allosteric modulator (basmisanil) failed to improve cognitive behaviour in DS^42^. This raises important questions regarding the over-inhibition hypothesis. While α5-GABA_A_Rs represent a small subset of receptors, the key issue is with the over-reliance on specific mouse models for probing mechanisms in DS. To date, the vast majority of findings pertaining to increased GABA signalling in DS, which has formed the centrepiece hypothesis for dysfunctional synaptic plasticity, stemmed from studies in Ts65Dn mice. However, the Ts65Dn strain also harbours an extra copy of 46 genes which are not orthologous to Hsa21^43^. This confounds the analysis of the Ts65Dn strain, since some phenotypes have been shown to be due to the increased dosage of these non-Hsa21 orthologous genes^44^.

To circumvent this concern, we investigated the DS GABA hypothesis using a different mouse model, Dp1Tyb. This strain has an extra copy of the entire region of mouse chromosome 16 (Mmu16) that is orthologous to Hsa21, including all of the Hsa21-orthologous genes that are increased in dosage in Ts65Dn mice but none of the irrelevant non-Hsa21 orthologous genes^45^. Thus, Dp1Tyb mice more closely reflect the increased gene dosage present in human DS with defects in learning, memory and neural activity^46,47^. They are genetically similar to Dp(16)1Yey mice which have the same number of Mmu16 genes in three copies and exhibit defective learning and memory, and impaired LTP^37,48,49^.

By using a more refined and relevant mouse model and studying three key brain areas for cognition, we show that, contrary to previous reports, over-inhibition is not an omnipresent feature of DS but presents as a brain region-specific phenomenon. Adult Dp1Tyb neurons feature increased GABA-mediated phasic and tonic inhibition in the hippocampal dentate gyrus (DG), whilst inhibition is unchanged in CA1 neurons where GABA acts as a classical inhibitory neurotransmitter. In the medial prefrontal cortex, layer II/ III neurons, there is also a negligible effect on phasic inhibition. Furthermore, using a genetic mapping panel we demonstrate that the increased tonic inhibition in the DG is caused by an extra copy of at least four genes and identify *Olig1*, *Olig2* and *Dyrk1a* as three of these causative genetic elements^51^.

## Results

### Selecting a DS mouse model

Hsa21 has homology to three regions in the mouse genome, located on Mmu10, Mmu16 and Mmu17. The largest region is on Mmu16 (Figure 1A; Supplementary Table 1). Dp1Tyb mice have an additional copy of the entire 144-gene Hsa21-orthologous region of Mmu16 and thus, genetically, model trisomy of around 63% of Hsa21^50^. The increased gene dosage in Dp1Tyb mice includes all of the Hsa21-orthologous genes that are in three copies in Ts65Dn mice, but none of the additional irrelevant genes, making it a more accurate DS model. Thus, we chose to use the Dp1Tyb strain to investigate the hypothesis that GABA-mediated inhibition is increased in the DS brain.

**Figure 1.**
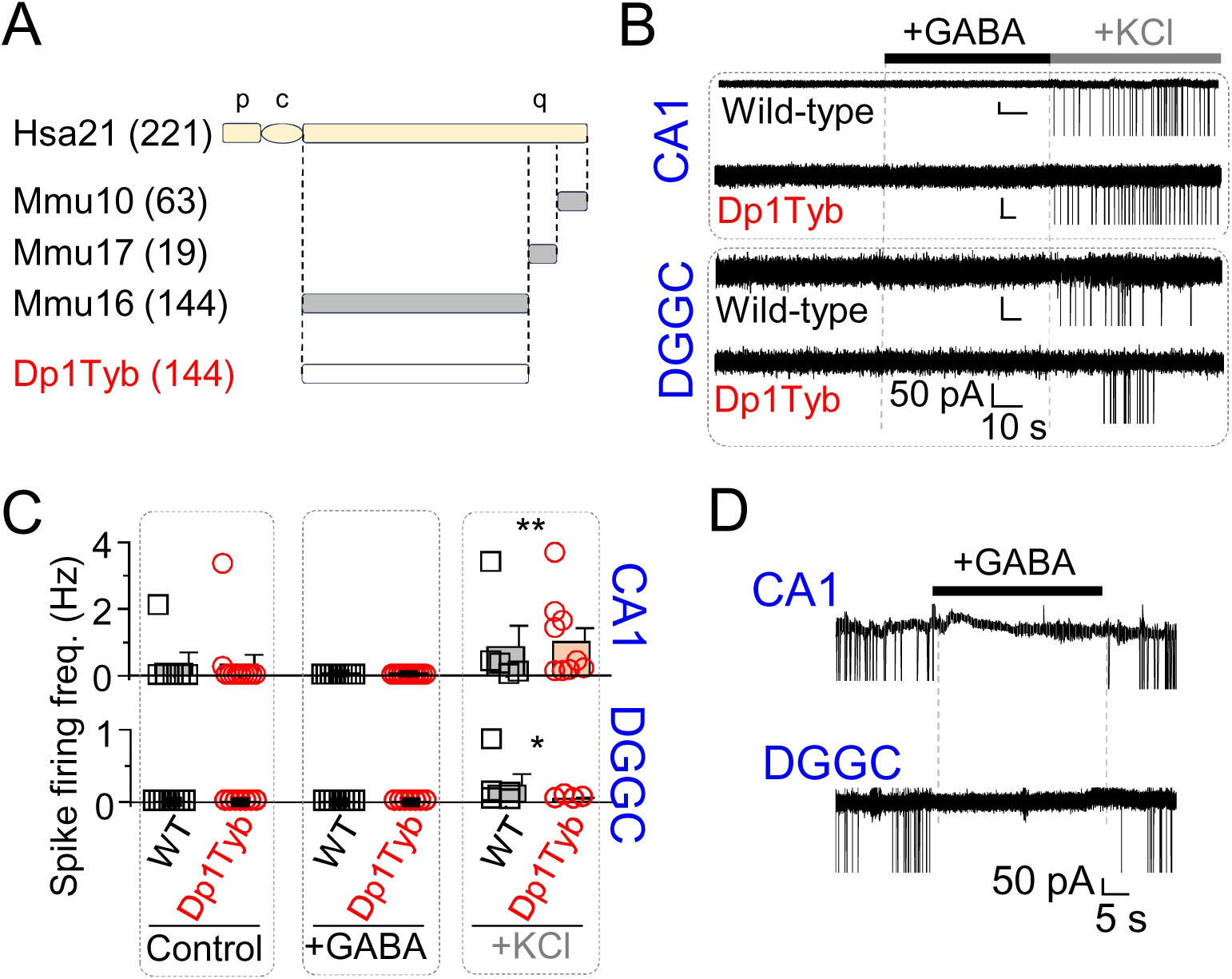
Inhibitory actions of GABA in acute brain slice hippocampal neurons. *A,* Schematic (not to scale) showing the short (p) and long (q) arms and the centromere (c) of Hsa21 (beige) aligned with orthologous regions on mouse chromosome (Mmu) 10, Mmu17 and Mmu16 (grey). The short arm of Hsa21 is highly repetitive with no obvious orthology to mouse chromosomes. Dp1Tyb mice contain a duplication of the entire Hsa21-orthologous region of Mmu16 (white), which is only a small part of the whole of Mmu16, the rest of which is not shown. Numbers in parentheses indicate numbers of coding genes in these intervals. *B,* Example cell attached recordings of CA1 and dentate gyrus granule cells (DGGCs) in control condition and in the presence of 100 μM GABA and 10 mM KCl from wild-type and Dp1Tyb mice. *C,* Spike firing frequency showing that action potentials are not induced by GABA but by KCl. *D,* Application of GABA blocks action potentials in spontaneously spiking DGGCs and CA1 neurons from Dp1Tyb mice. *P<0.05, **P<0.01 GABA compared to KCl, Kruskal-Wallis ANOVA, n = 6-12 cells and 3-4 animals.

### Region-specific change to GABA synaptic transmission in Dp1Tyb

To investigate the nature and extent of changes to GABA-mediated synaptic transmission, acute brain slices were prepared from adult (3-6 months) Dp1Tyb mice and their wild-type littermates as controls. Given prior observations that GABA can be excitatory in DS mice^40^ we first established whether this also featured in Dp1Tyb. The nature of GABA activity was ascertained using cell-attached recordings from neuronal somata to avoid disturbing the natural Cl^-^ electrochemical gradient. Notably, excitatory GABA activity, previously observed in some Ts65Dn studies^40^, was not evident in CA1 pyramidal neurons or in dentate gyrus granule cells (DGGCs) of the Dp1Tyb hippocampus. Bath application of 100 μM GABA did not generate action potentials (P>0.05, Kruskal-Wallis (KW) ANOVA) in wild-type or Dp1Tyb cells, whereas addition of 10 mM KCl to the same neurons, which causes depolarisation, initiated spike firing (Figure 1B-C; P<0.01, 0.05, KW ANOVA). Moreover, in spontaneously firing CA1 neurons and DGGCs, application of GABA inhibited action potentials that recovered on wash-off (Figure 1D). These results suggest that at this age, in Dp1Tyb neurons, GABA is operating as an inhibitory rather than excitatory transmitter, with the equilibrium potential for Cl^-^ (E_Cl_) presumably more negative than (or at least equal to) the resting membrane potential (E_m_).

The inhibitory profile of GABAergic signalling in Dp1Tyb was probed by whole-cell patch clamp recordings of neurons from three brain regions voltage clamped at −60 mV in the presence of 2 mM kynurenic acid to block excitatory synaptic transmission thereby facilitating the detection of GABA synaptic currents. The use of high internal Cl^-^ enabled the identification of spontaneous inhibitory postsynaptic currents (sIPSCs). Single cell recordings from CA1 pyramidal neurons in hippocampal slices of Dp1Tyb did not detect changes to whole cell capacitance, or averaged sIPSC amplitude and median frequency (compared to wild-type littermates (P>0.05, Figure 2A-B; Supplementary Figure 1A-B; Supplementary Table 2). We also analysed cumulative probability distributions for sIPSC amplitude and inter-event intervals but these remained unchanged in Dp1Tyb CA1 neurons compared to wild-type (P>0.05; Supplementary Figure 1A, C). By contrast, recordings from Dp1Tyb DGGCs revealed an increased sIPSC frequency (p = 0.0413, Mann-Whitney test), evident by the reduced inter-event interval (P<0.001; Mann-Whitney test), without affecting whole-cell capacitance, or the average or cumulative probability distributions of sIPSC amplitude and kinetics (P>0.05; Figure 2C-D; Supplementary Figure 1D-F, 2A-C). However, recordings of neurons in layers II/ III of the medial prefrontal cortex (mPFC) revealed unchanged (P>0.05) whole cell capacitance and unchanged averaged amplitude and frequency of sIPSCs signifying largely unchanged inhibition (Figure 2E-F; Supplementary Figure 1G-I). For both genotypes, in all three brain areas, 50 μM bicuculline blocked all synaptic activity indicating that the sIPSCs were mediated via synaptic GABA_A_ receptors (Supplementary Figure 1A, D, G).

**Figure 2.**
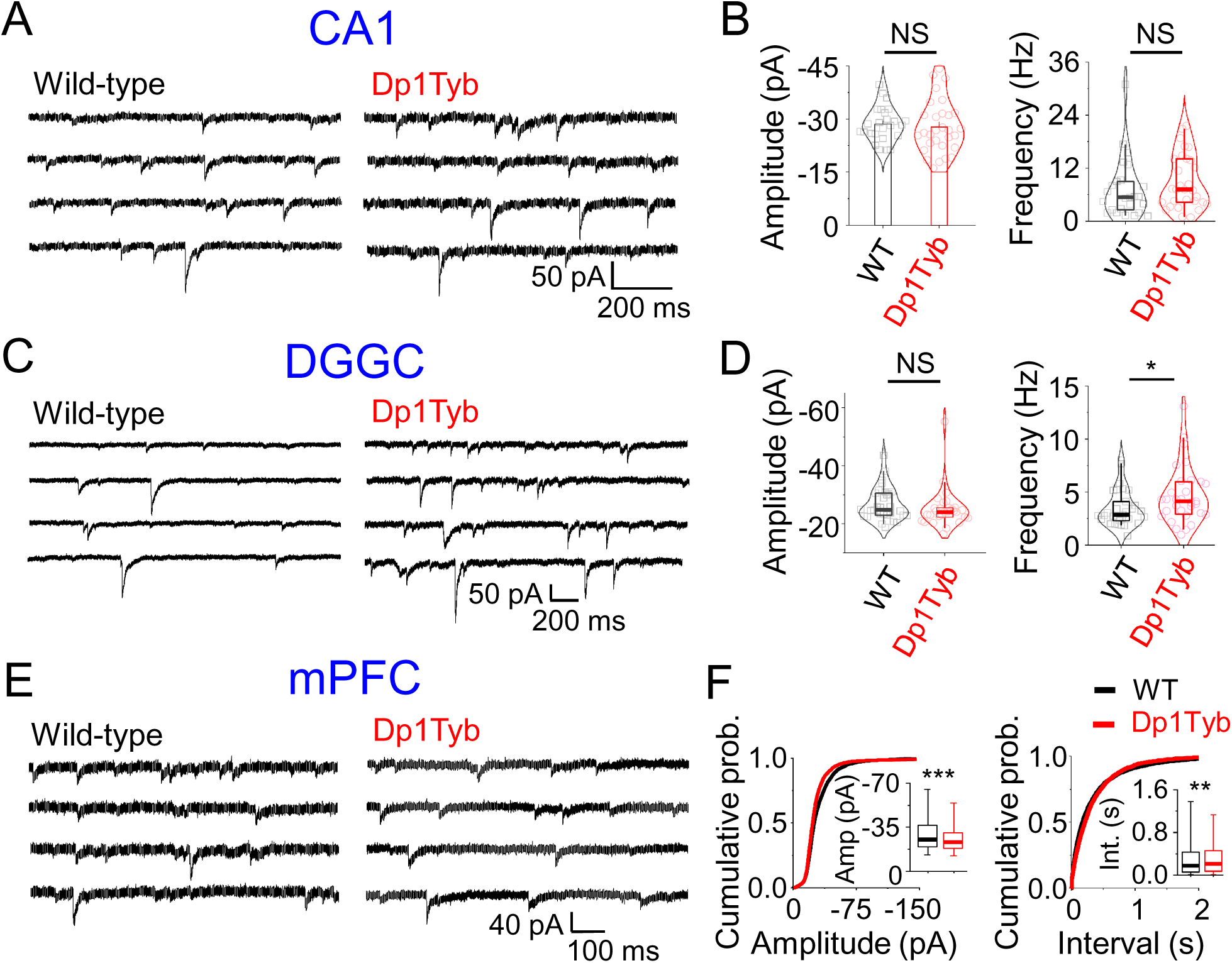
Brain area specific changes to phasic inhibition. *A,* Example spontaneous inhibitory postsynaptic current (sIPSC) recordings of CA1 pyramidal neurons from wild-type and Dp1Tyb mice. *B,* Unchanged sIPSC amplitude and frequency of CA1 neurons from Dp1Tyb compared to wild-type (WT). *C,* Representative sIPSC recordings of dentate gyrus granule cells (DGGCs) from wild-type and Dp1Tyb animals. *D,* Unchanged amplitude and increased sIPSC frequency of Dp1Tyb DGGCs compared to their wild-type counterparts. *E,* sIPSC recordings of layer II/ III medial prefrontal cortex (mPFC) neurons from wild-type and Dp1Tyb mice. *F,* Cumulative probability distributions of sIPSC amplitude and inter-event intervals of Dp1Tyb mPFC neurons compared to wild-type. Inset, boxplot showing median amplitude and interval. NS – not significant, *P<0.05, **P<0.01, ***P<0.001, two tailed unpaired t-test or Mann-Whitney test. n = 16-31 neurons, 7-9 animals.

The core hypothesis that GABA mediated synaptic inhibition is perturbed by trisomy, is therefore applicable to the Dp1Tyb DS model, but unexpectedly, this is critically dependent upon the brain region, with overall increased inhibition evident in DGGCs, but no overt change detected in the mPFC and hippocampal CA1 regions.

### Increased GABA-mediated tonic inhibition in the Dp1Tyb dentate gyrus

Granule cells of the DG exhibit limited spontaneous activity, being naturally restrained^51,52^ under the powerful control of tonic GABA inhibition^53^. To explore whether tonic inhibition was differentially regulated by increased gene dosage in Dp1Tyb, whole-cell patch clamp recording was used. The level of tonic current was measured as an outward current at −60 mV initiated by applying a saturating concentration of bicuculline (50 μM). Tonic current varied by up to 3-fold between brain regions in wild-type brain slices (Figure 3A-F). No change to the level of tonic current was apparent by recording from neurons in the CA1 hippocampal (p=0.9728, Mann-Whitney test) or mPFC (p=0.9227, two-tailed unpaired t-test) regions from Dp1Tyb slices. By contrast, in the dentate gyrus, tonic current was increased for Dp1Tyb compared to wild-type controls (p=0.0180, Mann-Whitney test). Thus, the only effect on tonic current was observed in the dentate gyrus, a location where tonic inhibition will most likely manipulate neural communication in this important brain area for cognition.

**Figure 3.**
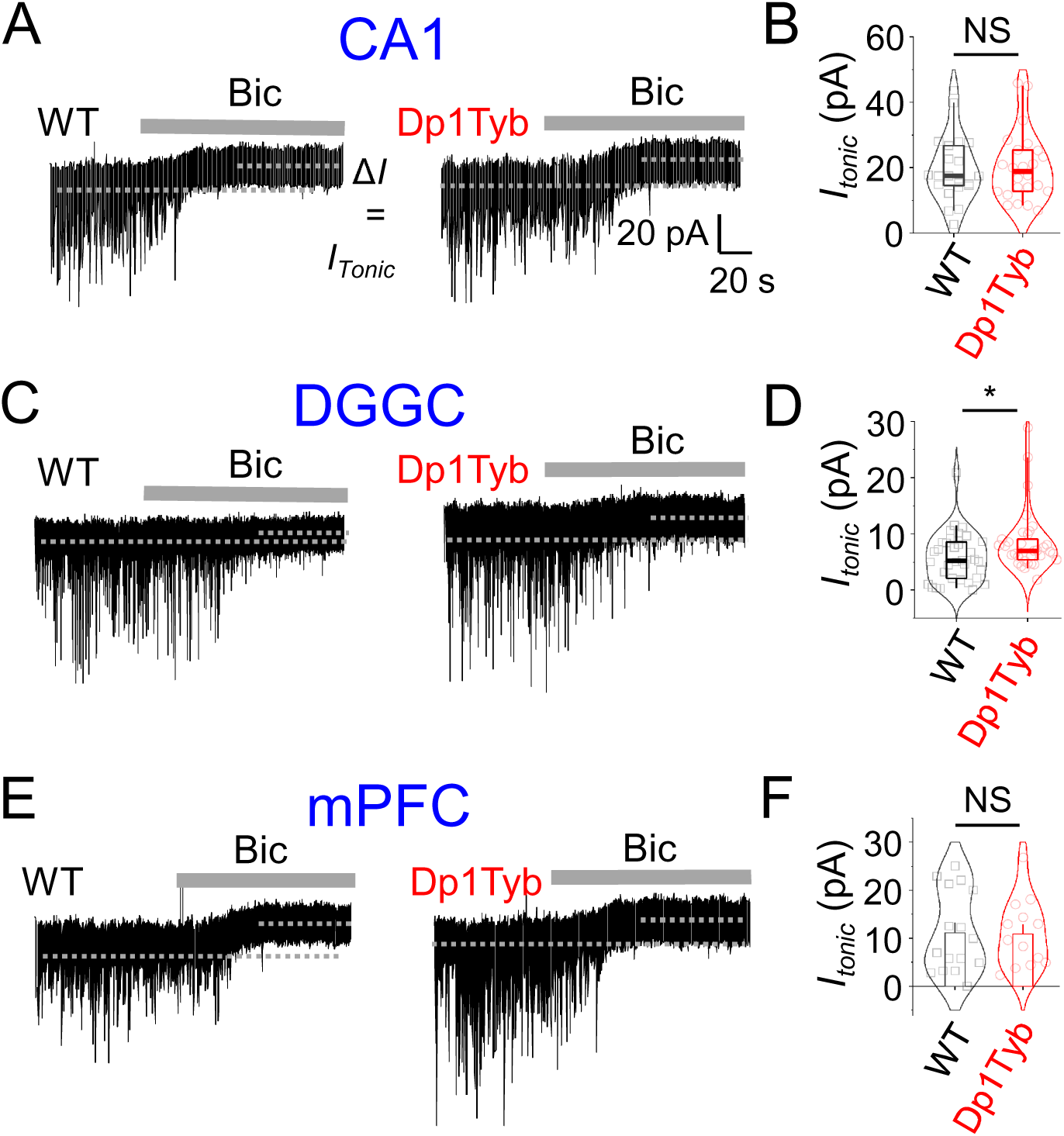
Brain area specific elevation of tonic inhibition. *A,* Example voltage clamp recordings of GABAergic events and tonic currents from wild-type (WT) and Dp1Tyb CA1 neurons. Grey lines depict application of 50 μM bicuculline (Bic) which abolishes inhibitory postsynaptic currents confirming their GABAergic nature. In addition, an upward defection of membrane current (difference between the dotted lines; Δ*I* = *I_Tonic_*) revels the extent of tonic inhibition. *B,* Unchanged tonic currents of Dp1Tyb CA1 neurons compared to wild-type. *C,* Representative GABAergic events and tonic current recordings of dentate gyrus granule cells (DGGCs) from wild-type and Dp1Tyb animals. *D,* Increased tonic inhibition of Dp1Tyb DGGCs compared to their wild-type counterparts. *E,* GABAergic events and tonic current recordings of layer II/ III medial prefrontal cortex (mPFC) neurons from wild-type and Dp1Tyb mice. *F,* Unchanged tonic inhibition of Dp1Tyb mPFC neurons compared to wild-type. NS – not significant, *P<0.05, Mann-Whitney test. n = 13-37 neurons, 7-9 animals.

### Altered neurophysiology of Dp1Tyb neurons subject to increased GABA inhibition

To assess the impact of enhanced GABA-mediated synaptic and tonic inhibition on DGGC neurophysiology, individual cells were recorded in whole-cell current clamp mode to assess their excitability. The mean resting membrane potential was similar between wild-type and Dp1Tyb neurons (P>0.05, two-tailed unpaired t-test; Figure 4A; Supplementary Table 3). Depolarising constant current steps were injected to measure action potential threshold (rheobase) and the number of spikes fired per current step. The rheobase was evidently higher (P<0.001, KW ANOVA) for Dp1Tyb cells and they supported a reduced spike firing frequency per depolarizing current step (Figure 4B-D; Supplementary Table 3) shifting the input-output curve without changes to the slope - an outcome that could arise from increased GABA-mediated tonic inhibition^54^. To establish if the spike rheobase could be reset to wild-type control levels, Dp1Tyb neurons were treated with 50 μM bicuculline. This reduced the rheobase (Figure 4D) and abolished (P>0.05, KW ANOVA) the shift in the input output curve (Figure 4E). Consistent with increased tonic inhibition, the membrane input resistance was reduced (P<0.01, One-way ANOVA) for Dp1Tyb DGGCs indicative of a persistent leak current which bicuculline normalised (P>0.05, One-way ANOVA; Figure 4F; Supplementary Table 3) towards that of wild-type neurons. By contrast, the spike firing properties of CA1 pyramidal neurons did not differ between wild-type and Dp1Tyb genotypes (Supplementary Figure 3A-E) and this likely reflects unchanged GABAergic inhibition or intrinsic membrane properties in this area. Similarly, in the absence of changes to tonic and phasic inhibition (Figure 2E-F) spike firing properties in layer II/ III neurons of the mPFC will not be influenced by changes to GABA inhibition in Dp1Tyb mice.

**Figure 4.**
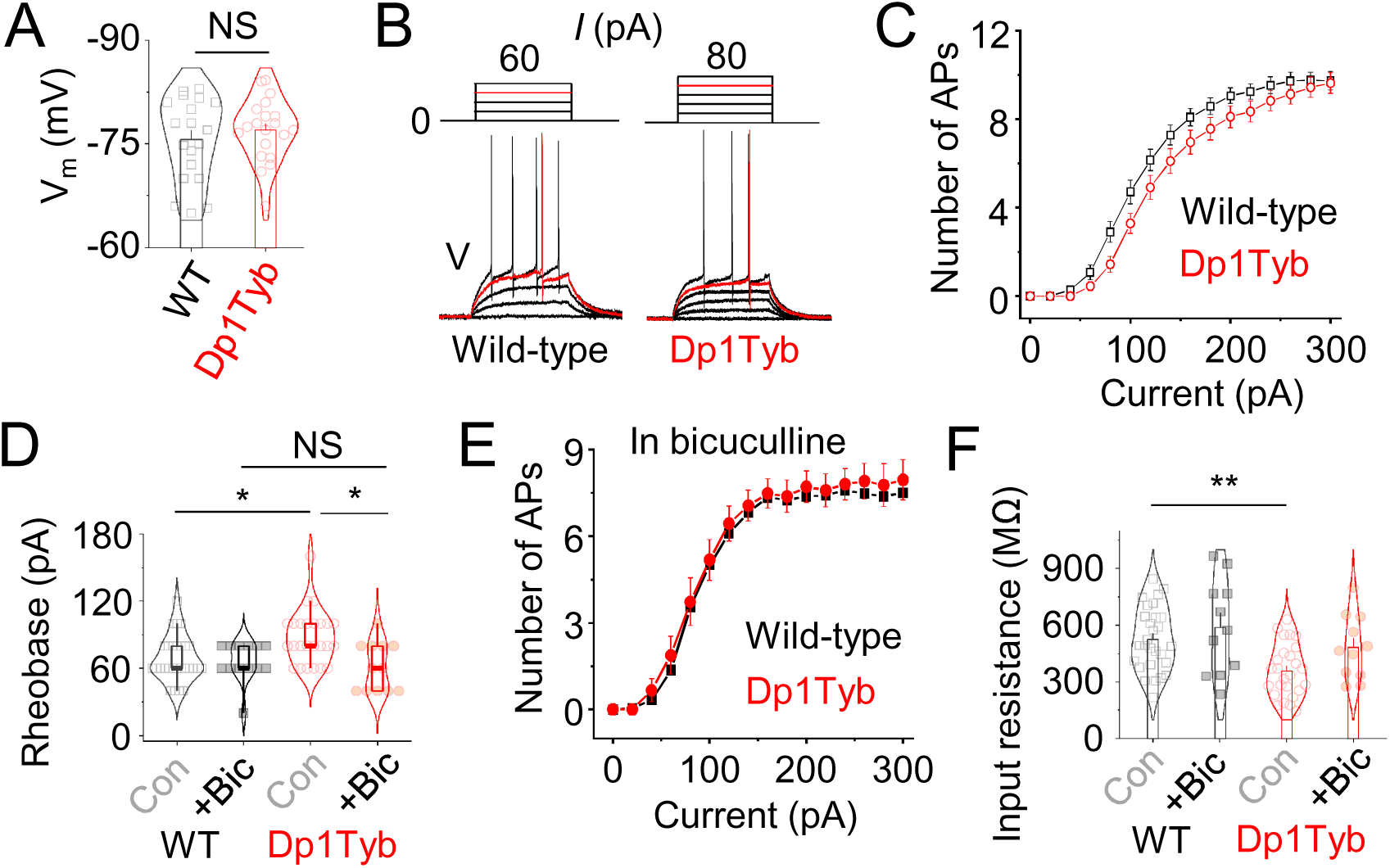
Elevated tonic inhibition alters spike firing properties in dentate cells. *A,* Average resting membrane potential of wild-type (WT) and Dp1Tyb dentate gyrus granule cells. *B,* Example current clamp recordings from wild-type and Dp1Tyb granule cells in response to constant step current injections. The rheobase trace has been depicted in red. *C,* Current-action potential relationship of wild-type and Dp1Tyb dentate gyrus granule cells. *D,* Rheobase of wild-type and Dp1Tyb granule cells in control and bicuculline (bic; 50 μM). *E,* Current-action potential relationship of dentate gyrus granule cells in bicuculline. *F,* Input resistance of wild-type and Dp1Tyb granule cells in control and bicuculline. NS – not significant, *P<0.05, **P<0.01, two-tailed unpaired t-test, One way-ANOVA or Kruskal-Wallis nonparametric ANOVA. n = 12-28 neurons, 4-5 animals.

These data indicate that an extra copy of one or more of the genes that are duplicated in Dp1Tyb mice is responsible for increased tonic inhibition in dentate gyrus neurons, and that this change results in the neurophysiological phenotype of reduced spike firing, and a higher threshold for spike initiation.

### Mechanisms underlying increased GABA inhibition in the dentate gyrus

The brain region-specific modulation of GABA inhibition, clearly exemplified by increased GABA-mediated tonic inhibition in the DG, could arise following alterations to several distinct signalling components affecting the GABA receptor activation pathway. These include: elevated extrasynaptic GABA_A_ receptor activity, remodelling of principal cell dendritic morphology, increased interneuron numbers and level of excitability, reduced activity of astrocytes and reduced synaptic GABA transporter capacity, or by increased presynaptic GABA release. All or some of these factors could be important for enhancing inhibition in Dp1Tyb neurons. The DG is unique in the hippocampus as the majority of tonic inhibition in granule cells is mediated by δ-subunit containing GABA_A_Rs^55^. These receptors can be selectively activated by the superagonist THIP (4,5,6,7-tetrahydroisoxazolo[5,4-c]pyridin-3-ol), at <1μM^56,57^. Application of 1 μM THIP to dentate granule cells in brain slices in the presence of tetrodotoxin (TTX, 0.5 μM) to prevent action potential driven release of GABA from interneurons, revealed larger THIP currents for Dp1Tyb neurons compared to their wild-type counterparts (Figure 5A-B; p = 0.0112, two-tailed unpaired t-test), suggesting that increased extrasynaptic δ-containing receptor numbers or their basal activity could underlie enhanced tonic GABA current in these neurons.

**Figure 5.**
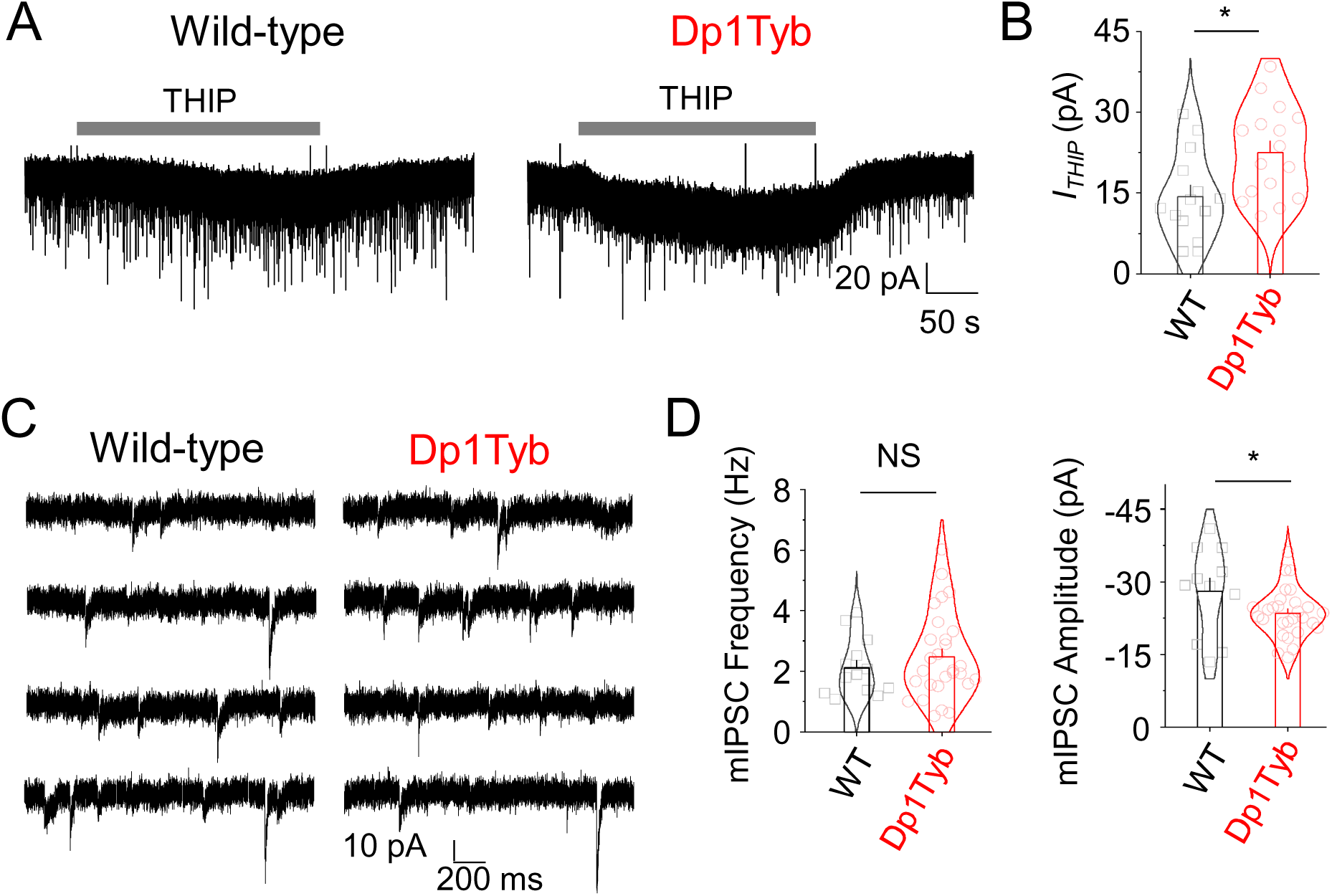
Elevated tonic inhibition via δ-GABA_A_Rs. *A,* Representative voltage clamp recordings of THIP (1 µM) currents from dentate granule cells in the presence of tetrodotoxin. *B,* Bar-graphs showing elevated THIP currents in Dp1Tyb neurons compared to wild-type (WT) cells in dentate gyrus. *C,* Representative traces of miniature inhibitory postsynaptic currents in the presence of tetrodotoxin of wild-type and Dp1Tyb dentate gyrus granule cells. *D,* mIPSC frequency and amplitude of granule cells. NS – not significant, *P<0.05, two-tailed unpaired t-test n = 14-28 cells; 6-8 animals.

In the presence of TTX, the frequency of miniature IPSCs (mIPSCs) was unaltered (P>0.05, two-tailed unpaired t-test) in Dp1Tyb DGGCs compared to wild-type equivalents, although an unexpected reduction of mIPSC amplitude was apparent (Figure 5C-D; p = 0.0239, two-tailed unpaired t-test). This suggested that the increased frequency of sIPSCs in Dp1Tyb DGGCs is action potential driven from increasingly excitable interneurons. To assess GABA inhibition at the synaptic level, peak-scaled non-stationary noise analysis of the IPSCs revealed no change in the estimated single channel conductance between Dp1Tyb and WT neurons (p=0.1057, two-tailed unpaired t-test), and no change (p=0.5819) in the number of synaptic receptors (Supplementary Figure 4A-C).

To investigate whether Dp1Tyb neurons exhibited altered morphology, particularly the dendritic arbour, granule neurons were filled with lucifer yellow via the patch electrode, followed by fixation, before performing a Sholl analysis on 3D reconstructed confocal images. This revealed increased dendritic arborisation, indicating principal cell remodelling for Dp1Tyb DGGCs (Figure 6A-C; P<0.05, two-tailed unpaired t-test). In addition, the number of immunolabelled GABA-positive (Figure 6D-F; p=0.0444, two-tailed unpaired t-test) and parvalbumin (PV)-positive (Figure 6G-I; p=0.0047, Mann-Whitney test) cells was also increased in the DG but not in CA1 (P>0.05), which could potentially increase GABAergic innervation and ambient GABA to support the overactive GABA phenotype that characterises DG neurons in Dp1Tyb slices.

**Figure 6.**
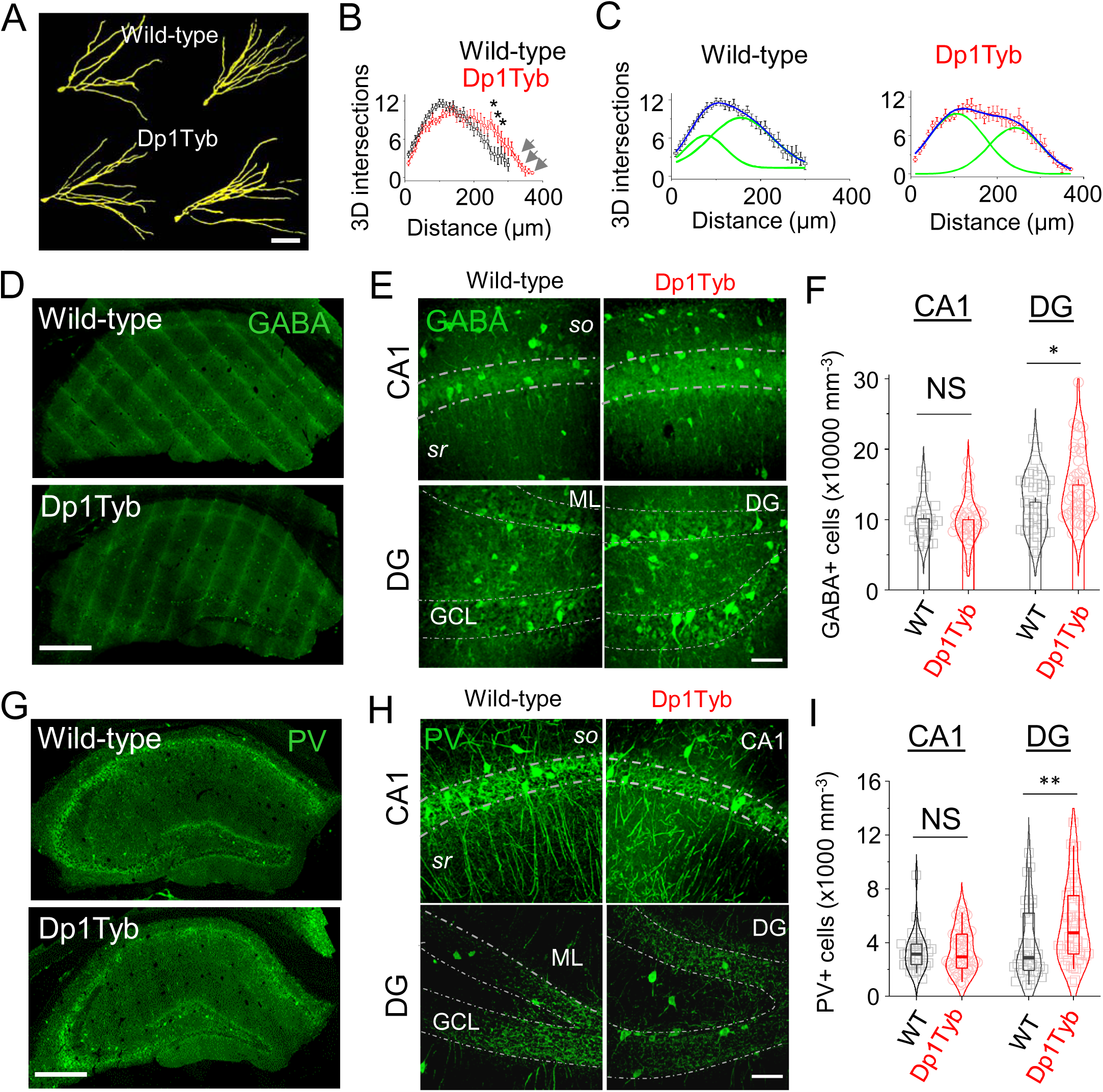
Altered dendritic morphology and interneuron numbers in dentate gyrus. *A,* Representative images of dye-filled wild-type (WT) and Dp1Tyb dentate gyrus granule cells. Scale bar 100 µm. *B,* 3D Sholl analysis of dendritic morphology of wild-type and Dp1Tyb neurons. Note – Dp1Tyb neurons extend further than their wild-type counterparts (grey arrows). *C,* Sholl analysis of dendritic morphology of wild-type and Dp1Tyb neurons fitted with two Gaussian components (green curves). *D,* Representative hippocampal GABA immunolabelling of wild-type and Dp1Tyb brain sections. Scale bar 500 µm. *E,* Images of GABA labelled CA1 and dentate gyrus (DG) neurons of wild-type and Dp1Tyb brain sections. *so*, *stratus oriens*; sr, *stratum radiatum*; ML, molecular layer; GCL, granule cell layer. Scale bar 50 µm. *F,* Bar graph showing the density of GABA interneurons in CA1 and DG sections. *G,* Representative hippocampal parvalbumin (PV) immunolabelling of wild-type and Dp1Tyb brain sections. Scale bar 500 µm. *H,* Images of PV labelled CA1 and DG interneurons of wild-type and Dp1Tyb brain sections. Scale bar 50 µm. *I,* Box plots showing the density of PV interneurons in CA1 and DG sections. *P<0.05, two-tailed unpaired t-test or Mann-Whitney test, n = 35-47 sections from, 5 animals per genotype.

Thus, our results indicate that increased GABA signalling in Dp1Tyb DGGC may be due to a combination of increased numbers or activity of extrasynaptic GABA_A_Rs, increased arborization of DGGC or increased numbers of interneurons including an increase in their excitability.

### Identifying genes causing the GABA neurophenotype in Dp1Tyb

While phasic inhibition controls, in a time-dependent manner, spike firing, bursting, and latency^58^, tonic inhibition determines the input-output gain by persistently increasing membrane conductance (causing shunting inhibition^59^). Phasic inhibition generally changes the slope of action potential firing input-output curves whereas tonic inhibition laterally shifts these curves along the current-axis^54^. The later was noted following analysis of spike-firing in Dp1Tyb DGGCs suggesting tonic is more influential in Dp1Tyb DGGCs which we subsequently focused on hereafter for gene mapping. To determine which of the 144 Hsa21 orthologous genes on Mmu16 are required in three copies to increase GABA signalling in Dp1Tyb DGGCs we utilised our panel of 8 chromosome-engineered mouse lines (Dp2Tyb - Dp9Tyb). Each of these lines duplicate a smaller genomic region of Mmu16, which are within the complete region duplicated in Dp1Tyb mice (Supplementary Table 1)^50^. When summed these small regions cover the entire region that is increased in copy number in the Dp1Tyb line.

Initially, we analysed Dp9Tyb, Dp2Tyb and Dp3Tyb mice which retain extra copies of 72, 33 and 39 genes, respectively (Figure 7A; Supplementary Table 1). GABA transmission was assessed in the DG of acute slices by single cell patch clamp electrophysiology. Both Dp2Tyb and Dp3Tyb mice exhibited an enhanced level of tonic and phasic inhibition (Figure 7B-C; Supplementary Table 2). In contrast, tonic current was unchanged in Dp9Tyb DGGCs compared to wild-type neurons (Figure 7D). This suggests that increased dosage of more than one gene is responsible for the GABA phenotype observed in Dp1Tyb mice, and that there is at least one causative dosage-sensitive gene located in each of two distinct genomic regions, that are present in three copies in Dp2Tyb and Dp3Tyb mice.

**Figure 7.**
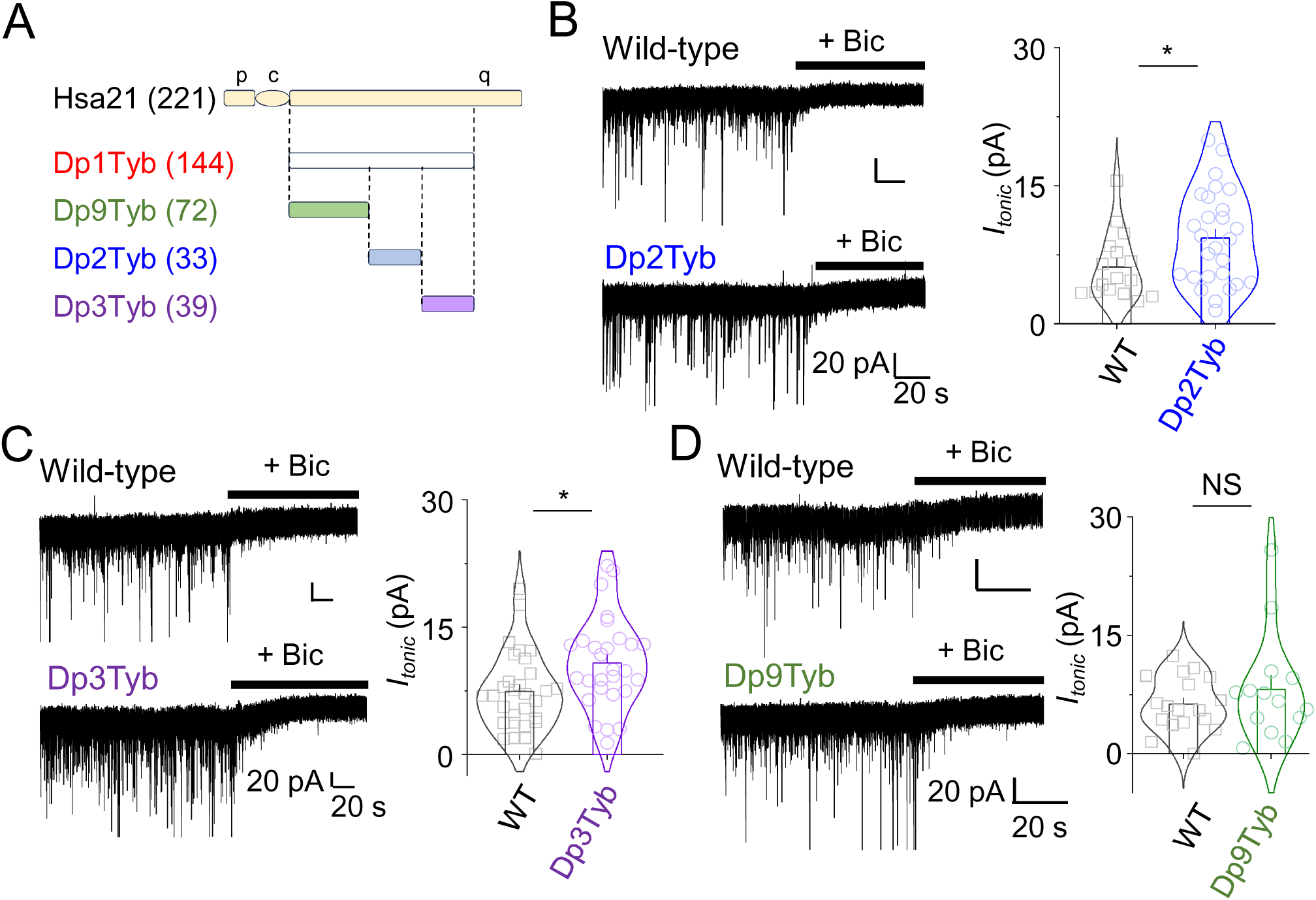
Elevated tonic inhibition due to duplication of two distinct regions of Mmu16. *A,* Schematic showing the short (p) and long (q) arms and the centromere (c) of Hsa21 aligned with orthologous regions on mouse chromosome 16 (Mmu16) that are present in a third copy in Dp1Tyb, Dp2Tyb, Dp3Tyb and Dp9Tyb mice. Numbers in parentheses indicate numbers of coding genes in these intervals. *B,* Example voltage clamp recordings of GABAergic events and tonic current from wild-type (WT) and Dp2Tyb dentate gyrus granule cells along with average tonic currents . Dark horizontal lines depict application of bicuculline (+bic; 50 μM) which abolishes inhibitory postsynaptic currents confirming their GABAergic nature. *C,* Representative tonic current traces of WT and Dp3Tyb dentate granule cells along with average tonic currents. *D,* Example voltage clamp traces of tonic currents from WT and Dp9Tyb dentate gyrus granule neurons along with average tonic currents. NS – not significant, *P<0.05, two-tailed unpaired t-test. n = 14-30 cells; 6-9 animals.

To further narrow down the location of the causative genes, we made use of mouse strains Dp7Tyb and Dp8Tyb which subdivide the Dp2Tyb region, and Dp4Tyb, Dp5Tyb and Dp6Tyb which subdivide the Dp3Tyb region (Supplementary Table 1; Supplementary Figure 5A)^50^. Analysis of GABA inhibition in the dentate gyrus of acute hippocampal slices from each of these 5 mouse strains compared to wild-type controls revealed no change in tonic GABA current or in phasic inhibition (Supplementary Figure 5B-F; Supplementary Table 2). This loss of the GABA phenotype in the smaller duplication strains implies that there must be at least two causative genes in each of the Dp2Tyb and Dp3Tyb regions, i.e. a total of at least 4 genes.

### Increased dosage of *Olig1*, *Olig2* and *Dyrk1a* cause increased tonic inhibition in DG

To identify specific genes whose increased dosage in DS may cause the GABA phenotype, we tested the potential involvement of three candidate genes: *Olig1*, *Olig2* and *Dyrk1a*. *Olig1* and *Olig2* are located next to each other within the region that is duplicated in Dp2Tyb and Dp7Tyb mice and encode transcription factors which have been implicated in neurogenesis, including that of interneurons (Figure 8A)^60,61^. We note that, reducing the copy number of *Olig1* and *Olig2* from 3 to 2 in Ts65Dn mice reversed the increased generation of interneurons^23^. *Dyrk1a* is located in a region that is duplicated in Dp3Tyb and Dp5Tyb mice and encodes a kinase whose overexpression has been linked to elevated differentiation of GABAergic neurons and increased inhibition (Figure 8A)^62–64^. To test the involvement of these genes in the increased GABA-mediated tonic inhibition, we crossed mice with a heterozygous mutation of both *Olig1* and *Olig2* (*Olig1*^+/-^*Olig2*^+/-^) to Dp2Tyb mice to generate Dp2Tyb*Olig1*^+/+/-^*Olig2*^+/+/-^ (Dp2Tyb*Olig1/2*KO) mice in which the copy number of the *Olig1* and *Olig2* genes was reduced from 3 to 2 (Figure 8B). Similarly, we crossed mice with a heterozygous loss of function of *Dyrk1a* (*Dyrk1a*^+/-^) with Dp3Tyb to generate Dp3Tyb*Dyrk1a*^+/+/-^ (Dp3Tyb*Dyrk1a*KO) mice with the copy number of *Dyrk1a* reduced from 3 to 2 (Figure 8B).

**Figure 8.**
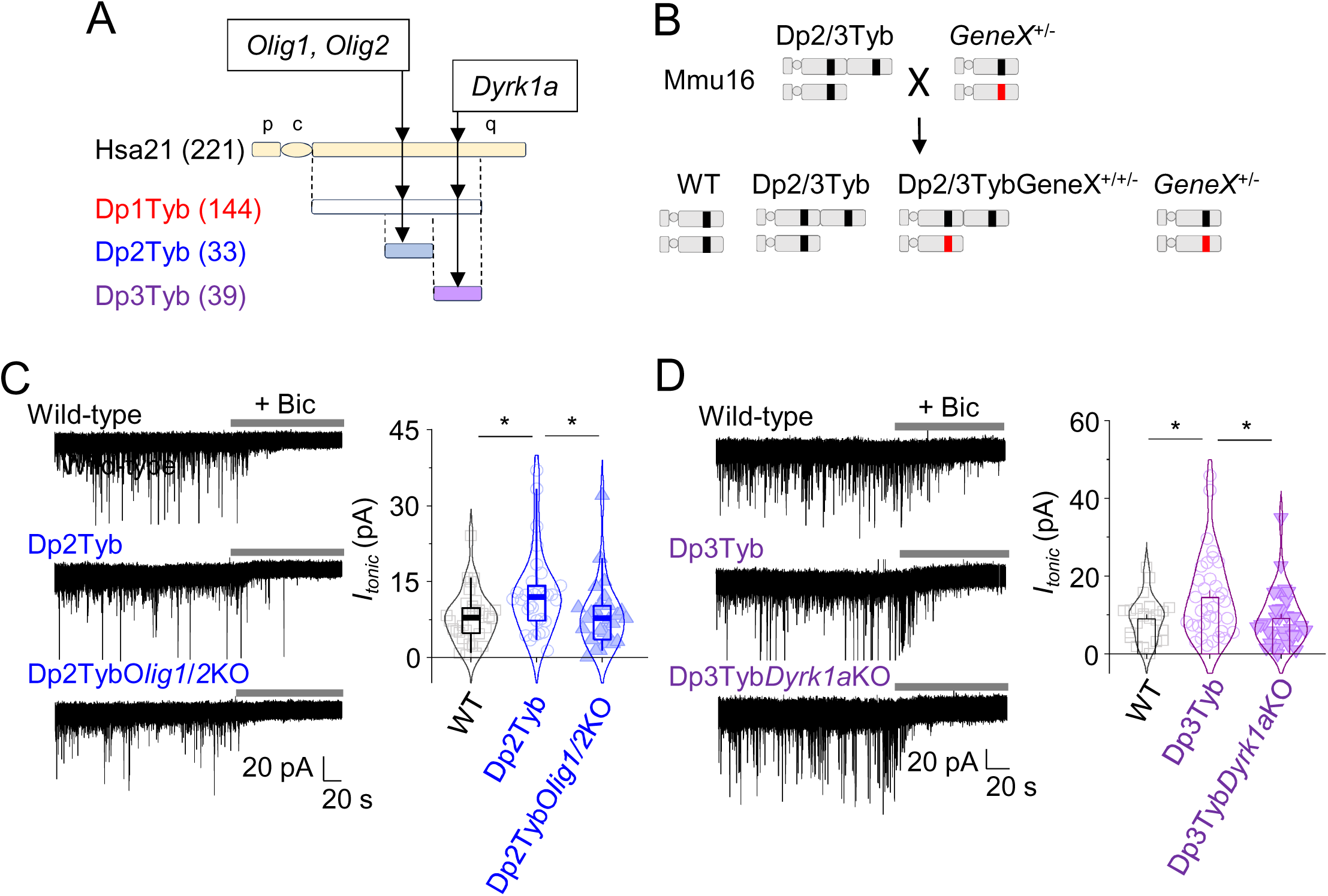
Tonic inhibition rescue by reduction of *Dyrk1a* and *Olig1*-2 copy numbers. *A,* Schematic showing the short (p) and long (q) arms and the centromere (c) of Hsa21 aligned with orthologous regions on mouse chromosome 16 (Mmu16) that are present in a third copy in Dp1Tyb, Dp2Tyb and Dp3Tyb mice, indicating the location of the *Olig1*, *Olig2* and *Dyrk1a* genes that are present in three copies in Dp1Tyb mice and in either Dp2Tyb or Dp3Tyb mice. *Olig1* and *Olig2* are adjacent to each other in the genome. The orthologous human genes (*OLIG1*, *OLIG2*, *DYRK1A*) are located in the equivalent locations on Hsa21. Numbers in parentheses indicate numbers of coding genes in these intervals. *B,* Pictogram showing genetic strategy used to reduce copy numbers of candidate genes from three to two in Dp2Tyb or Dp3Tyb mice. Crossing Dp2Tyb or Dp3Tyb mice (Dp2/3Tyb) carrying three copies of the candidate gene (black line) to strains with a heterozygous loss of function allele of either *Olig1* and *Olig2* or *Dyrk1a* (*GeneX*^+/-^, red line) results in progeny that include wild-type mice, Dp2Tyb or Dp3Tyb mice, and Dp2Tyb or Dp3Tyb mice with just two functional copies of GeneX (Dp2/3Tyb*GeneX*^+/+/-^). *C,* Voltage clamp recordings and GABAergic tonic currents from wild-type (WT), Dp2Tyb and Dp2Tyb*Olig1*^+/+/-^*Olig2*^+/+/-^ (Dp2Tyb*Olig1/2*KO) dentate gyrus granule cells; in the latter strain, the copy number of *Olig1* and *Olig2* has been reduced from 3 to 2. *D,* Voltage clamp recordings and GABAergic tonic currents from wild-type, Dp3Tyb and Dp3Tyb*Dyrk1a*^+/+/-^ (Dp3Tyb*Dyrk1a*KO) dentate gyrus granule cells; in the latter strain, the copy number of *Dyrk1a* has been reduced from 3 to 2. *P<0.05, One way-ANOVA or Kruskal-Wallis nonparametric ANOVA. n = 23-34 cells, 5-10 animals.

Preparing acute brain slices and whole cell recording from DGGCs revealed that the dual reduction in copy number for *Olig1* and *Olig2* from 3 to 2 in the Dp2Tyb animals, prevented the increase in tonic current (Figure 8C, Supplementary Table 4). Similarly, tonic current was also decreased to wild-type levels by reducing the copy number of *Dyrk1A* from 3 to 2 in Dp3Tyb mice (Figure 8D). Thus, increased dosage of *Olig1* and *Olig2* in Dp2Tyb mice and of *Dyrk1a* in Dp3Tyb mice is required, in part, for the increased GABA-mediated tonic inhibition.

## Discussion

Previous studies of neuronal signalling in DS, based exclusively on mouse models, have proposed that the condition is characterized by an excess level of GABA-mediated inhibition. However, premature termination of a clinical trial based on inhibiting α5-GABA_A_Rs casts doubt over this central tenet. In mitigation, there could be issues related to the drug efficacy and whether targeting α5-GABA_A_Rs for improving cognition in DS was appropriate; also we need to factor in the use of mouse models that overexpress genes in the nervous system that are unrelated to DS. Taken together, it seems a comprehensive re-evaluation of the role of GABA in DS is both timely and warranted.

To address this, we chose to investigate GABA signalling in the Dp1Tyb mouse model, which has an extra copy of the entire 144-gene Hsa21-orthologous region of Mmu16, including all the Hsa21-orthologous genes found in Ts65Dn mice, but not including additional irrelevant genes^50^. Furthermore, Dp1Tyb mice have learning and memory deficits, and altered neuronal activity^46,47^. In addition, we had previously generated an associated panel of mouse strains with shorter duplications, which allow the location of causative genes to be mapped^50,65^. Our study with Dp1Tyb mice suggests that excessive inhibition is brain circuit region-specific, with greater importance placed on changes to tonic rather than phasic inhibition. Moreover, we propose that duplication of at least 4 discrete genes underpins the GABA neurophenotype in DS and that targeting δ-GABA_A_Rs with improved mouse models may form a tractable therapeutic target for DS.

### Nature of GABA neurotransmission in Dp1Tyb mice

Several key discoveries emerged from our analysis of Dp1Tyb mice. Firstly, GABA neurotransmission was overtly inhibitory^41^ rather than excitatory^40^, in contrast to the conclusions of a previous study in Ts65Dn mice. The reasons for this difference are unclear but may involve overactivity of the Cl^-^ transporter NKCC1^67^ in Ts65Dn mice, potentially caused by increased dosage of one of the non-Hsa21 orthologous genes that are important for neurodevelopment and function^66^. Thus, it is impossible to know if neural phenotypes in Ts65Dn mice are due to an increased dosage of Hsa21-orthologous genes or the non-Hsa21 orthologous genes^44^.

The second notable observation is that the changes to GABA neurotransmission are not global throughout the Dp1Tyb brain but appear to be regional with inhibition in the DG featuring increased sIPSC frequency without change to the mean amplitude, whilst in the mPFC, and in the CA1 hippocampus, no overt changes to inhibition were evident. In addition to GABA synaptic inhibition, tonic inhibition, also mediated by GABA, was increased in the DG but not in the CA1 or mPFC regions. The third important feature of GABA inhibition in the Dp1Tyb model was revealed by probing the synaptic and tonic currents. Comparing miniature IPSCs to spontaneous IPSCs indicated that only sIPSCs showed an increased frequency in Dp1Tyb mice indicating that action potential-driven events underpinned the frequency change; indeed, mIPSCs showed no change in frequency and a small reduction in amplitude. At a molecular level, using non-stationary noise analysis of synaptic currents, suggested that single channel conductance for synaptic GABA_A_ receptors was unaffected by trisomy-21, and the estimated synaptic receptor number although trending to lower values remained statistically unchanged. The reason for the reduced mIPSC amplitude remains unclear but may involve signal filtering via the more extended dendritic arbour of Dp1Tyb DGGCs.

By contrast, the raised tonic inhibition in the DG was potentially due to increased δ-subunit containing GABA_A_ receptors or to a change in their activity, and this was reflected by the increased responses to the GABA_A_ receptor superagonist THIP, used at concentrations that selectively activate δ subunit-containing extrasynaptic receptors^57,67^. The dominant extrasynaptic GABA_A_ receptor in the DG expected to carry most of the tonic current is α4β3δ^68,69^, supplemented with a mix of α1-3β2/3γ2 receptors transitioning to and from inhibitory synapses^12,70,71^. Therefore, our results may explain why a phase II trial^73^ with the α5 negative allosteric modulator, basmisanil, projected to reduce tonic and perisynaptic GABA inhibition, failed to improve cognitive performance in DS volunteers^42^. This outcome may reflect that basmisanil was not targeting α4β3δ receptors, the dominant population in this region supporting increased tonic inhibition.

### Single cell and circuit network level changes in the Dp1Tyb model

At a single cell level, under current clamp conditions, native cell excitability changes occur in the Dp1Tyb DG that are in accord with increased GABA inhibitory tone. For instance, the increased spike firing threshold (increased rheobase), reduced spike frequency following the injection of a depolarising constant current step and reduction in the membrane input resistance are all expected in the presence of increased GABA_A_ receptor activity. All these changes in Dp1Tyb were reversed to control levels by addition of the GABA_A_ receptor antagonist bicuculline, confirming the increased GABA inhibition phenotype of this DS mouse line.

At a network level, there are morphological and cell population changes, including increased dendritic arborisation of Dp1Tyb DG granule cells coupled with increased numbers of GABAergic cells and parvalbumin-expressing interneurons, all of which would be commensurate with increased GABA neurotransmission in the Dp1Tyb DG.

At the neural circuit level, neural activity within the mPFC and the hippocampus, including their direct and indirect connectivity, is important for cognitive function^72^. At the synaptic level, GABA inhibition is largely unaltered in the mPFC whilst an increase is observed in the hippocampal DG region. Such changes to GABA inhibition are likely to impact on, and may underlie, several features of cognition. Local field potential and EEG recordings of Dp1Tyb mice have been coupled to a spatial working memory behavioural test - the spontaneous alternation T-maze^36^. This test is designed to distinguish between choosing the correct arm of the T-maze to explore and the decision-making time required to make this choice (latency). For Dp1Tyb mice, several aberrant features of behaviour were reported^46^. Dp1Tyb mice displayed greater immobility and slower decision making compared to wild-type control mice. Moreover, theta rhythm was reduced in the mPFC and hippocampus whilst theta-high gamma rhythm coupling was increased, which may be related to altered GABA transmission. Overall, whilst the Dp1Tyb mice took longer to decide which arm to enter, they invariably still made the correct choice. The dentate gyrus is regarded as a key area for memory and learning. It is the gateway to the hippocampus and will affect the overall functional output of this key brain structure. Interestingly, in Dp(16)1Yey mice, which have an additional copy of the same 144 genes as Dp1Tyb mice, spike generation profiles of CA1 pyramidal neurons are altered, potentially as a consequence of changes in GABA inhibition in the adjacent DG^73,74^. In summary, our findings suggest that the hypothesis of a global change in GABA-mediated inhibition across the DS brain is unlikely to be true, arguing instead in support of local changes to brain circuits.

### Mechanisms driving increased GABA inhibition in the DG

Our study of increased GABA inhibition in Dp1Tyb suggests multiple mechanisms are involved. We observed an increase in parvalbumin interneuron numbers, principal cell dendrite remodelling in the DG granule cells, and an increase in GABA tonic current potentially linked to the GABA sensitivity of extrasynaptic GABA_A_ receptors. However, notable questions remain, for example, what is the source of ambient GABA? We expected the increased frequency of sIPSCs might provide sufficient spillover GABA to activate extrasynaptic receptors but tracking GABA inhibition with our gene mapping panel suggest that the source of elevated tonic inhibition is likely to reside elsewhere. In this regard, in addition to extrasynaptic receptor activity, potential dysfunction of GABA transporters in neurons as well as astrocytes, could be an interesting area for future studies. In addition, our NSNA suggests that synaptic GABA_A_Rs in Dp1Tyb DGGCs are not impaired and thus raised tonic inhibition is the dominant change in GABA inhibition potentially facilitated by increased excitability of interneurons.

### Multiple genes contribute to the increased tonic GABA signalling in Dp1Tyb

Analysis of GABA signalling in the panel of mouse strains, each line containing a third copy of shorter variable length regions revealed that both Dp2Tyb and Dp3Tyb lines had increased GABA tonic inhibition in the DG. In contrast, analysis of the Dp7Tyb and Dp8Tyb strains that subdivide the Dp2Tyb region, and Dp4Tyb, Dp5Tyb and Dp6Tyb that subdivide the Dp3Tyb region, showed that none of these lines had altered GABA inhibition. Collectively, these results imply that there must be two or more genes in the 33-gene region that is duplicated in Dp2Tyb mice causing increased tonic inhibition, with at least one gene in each of the Dp7Tyb and Dp8Tyb regions. Separately, there must be two or more causative genes in the 39-gene region that is duplicated in Dp3Tyb, with at least one gene in each of at least two of the Dp4Tyb, Dp5Tyb or Dp6Tyb regions. Thus, in total we propose there are at least 4 causative genes. Given that Dp2Tyb and Dp3Tyb lines both exhibit increased tonic inhibition and have different non-overlapping sets of genes in three copies, the underlying pathological mechanisms in these two regions may be different.

By examining potential contributions to GABA inhibition made by individual candidate genes, we showed that reducing the dosage of *Olig1* and *Olig2* in Dp2Tyb mice or of *Dyrk1a* in Dp3Tyb mice, part-rescued the tonic inhibition phenotype. In the case of the *Olig* genes, we cannot tell whether increased dosage of *Olig1* or *Olig2*, or both, is required for the GABA inhibition phenotype, since the genes are closely linked and we used a mutant allele that inactivates both genes. The OLIG1 and OLIG2 transcription factors and the DYRK1A kinase may play a role in the generation of interneurons ^60–64^. Thus, their increased expression in Dp1Tyb mice may result in elevated tonic inhibition as a consequence of increased interneuron numbers.

Thus, in conclusion, we identified several regions on Mmu16 that underpin the GABA phenotype, showing that 4 or more genes are likely to contribute.

We also demonstrate that increased dosage of the *Olig1* and *Olig2* transcription factors and the *Dyrk1a* kinase contribute to the increased GABA inhibition and that deletion of just one copy of the transcription factors, or of the ubiquitous kinase, without further adjustment to the dosage of other potential contributory genes, is sufficient to normalise GABA activity in the Dp1Tyb DS model.

## Materials and Methods

Mice carrying the Dp(16Lipi-Zbtb21)1TybEmcf (Dp1Tyb), Dp(16Mis18a-Runx1)2TybEmcf (Dp2Tyb), Dp(16Mir802-Zbtb21)3TybEmcf (Dp3Tyb), Dp(16Mir802-Dscr3)4TybEmcf (Dp4Tyb), Dp(16Dyrk1a-B3galt5)5TybEmcf (Dp5Tyb), Dp(16Igsf5-Zbtb21)6TybEmcf (Dp6Tyb), Dp(16Lipi-Hunk)9TybEmcf (Dp9Tyb), *Olig1*^tm1And^ (*Olig1*^-^), *Olig2*^tm1And^ (*Olig2*^-^) and *Dyrk1a*^tm1Mla^ (*Dyrk1a*^-^) alleles have been described previously^50,75,76^. The generation of mice carrying Dp(16Mis18a-Il10rb)7TybEmcf (Dp7Tyb) and Dp(16Ifnar1-Runx1)8TybEmcf alleles will be described elsewhere (EL-E, SW-S, EMCF and VLJT, in preparation). The *Olig1*^tm1And^ and *Olig2*^tm1And^ alleles were linked in adjacent genes and hence transmitted together. To test candidate genes, we crossed Dp2Tyb mice to *Olig1*^+/-^*Olig2*^+/-^ mice to generate Dp2Tyb*Olig1*^+/+/-^*Olig2*^+/+/-^ (Dp2TybOlig1/2KO) mice containing two wild-type and one disrupted allele of each of *Olig1* and *Olig2*. Similarly, Dp3Tyb mice were crossed to *Dyrk1a*^+/-^ mice to generate Dp3Tyb*Dyrk1a*^+/+/-^ (Dp3TybDyrk1aKO) mice. All mice were bred and maintained on a C57BL/6J background (backcrossed for ≥ 10 generations) at the Francis Crick Institute and transferred to University College London where they were housed in specific-pathogen free conditions and given water and food ad libitum. Genotyping was performed using custom probes (Transnetyx). Mice were used at 3-6 months of age. Both male and female mice were used, and in all experiments sex and age were matched between mutant and control mice. Animal experiments were carried out under the authority of Project Licences granted by the UK Home Office and were approved by the Animal Welfare Ethical Review Bodies of the Francis Crick Institute and University College London. Numbers of protein-coding genes in different mouse strains were determined using the Biomart function in Ensembl on mouse genome assembly GRCm39, filtering for protein-coding genes, excluding three genes: ENSMUSG00000116933 which is a partial transcript for *Atp5o* (ENSMUSG00000022956), *Gm49711*, which is an alternatively spliced form of *Mrps6*, and *Gm49948* which is a fusion transcript of some exons from *Igsf5* and *Pcp4* (Supplementary Table 1). Note that the numbers of duplicated coding genes in the Dp strains have changed since our original publication^50^, and are likely to change further due to refinement of gene annotation. Similarly, using Biomart on human genome assembly GRCh38 identified 221 protein-coding genes on Hsa21.

### Preparation of brain slices

Acute brain slices were prepared from adult (3–6-month-old) mice in accordance with the UK Animals (Scientific Procedures) Act 1986. Brains collected after terminal anaesthesia with isoflurane were immersed in ice-cold slicing solution continuously bubbled with 95% air and 5% CO_2_ as described previously^77^. For the dentate gyrus, this was composed of (mM): 85 NaCl, 2.5 KCl, 1 CaCl_2_, 4 MgCl_2_, 1.25 NaH_2_PO4, 26 NaHCO_3_, 75 sucrose, and 25 glucose, pH – 7.4; for CA1 pyramidal and medial prefrontal cortex (mPFC) neurons a K gluconate-based slicing solution was used composed of (mM): 130 K-gluconate,15 KCl, 0.05 EGTA, 20 HEPES, 4 Na-pyruvate, 25 glucose, pH to 7.4 with KOH. Transverse 250 μm slices for dentate gyrus and coronal 350 μm slices for CA1/ mPFC were cut with a Leica VT1200S vibroslicer. The slicing solution was exchanged at 37 °C for 60 min with a recording solution containing (mM): 125 NaCl, 2.5 KCl, 2 CaCl_2_, 1 MgCl_2_, 1.25 NaH_2_PO4, 26 NaHCO_3_, and 25 glucose, pH 7.4. For recording inhibitory postsynaptic currents (IPSCs), 2 mM kynurenic acid was added to the slicing and recording solutions.

### Electrophysiology

Whole cell recordings were carried out after equilibrating slices at room temperature for 1 hr using an Axopatch 200B amplifier (Molecular Devices). Thin-walled borosilicate glass capillaries (GC150TF; World precision instruments) of 3–5 MΩ resistance filled with an internal solution^78^ containing (mM): 140 CsCl, 2 NaCl, 2 MgCl_2_, 5 EGTA, 30 KOH, 10 HEPES, 0.5 CaCl_2_, and 2 Na_-_ATP, 0.5 Na-GTP, 2 QX-314; pH – 7.3 with CsOH for recording IPSCs. For current clamp recording, the internal solution^79^ comprised (in mM): K-gluconate (130), NaCl (4), K-HEPES (10), K-EGTA (5), CaCl_2_ (0.5), Mg-ATP (2), pH to 7.3 with KOH. IPSC recordings were carried out in the presence of 2 mM kynurenic acid and current clamp recordings were carried out in the absence of any blockers unless otherwise stated.

Membrane currents were filtered at 5 kHz (−3 dB, 6th pole Bessel, 36 dB/ octave) and cells were voltage clamped at −60 mV with optimised series resistance (Rs, <10 MΩ) and whole-cell membrane capacitance compensation. For current clamp recordings, neurons were held at resting membrane potentials with zero current injection and the resting membrane potential of neurons were noted immediately after establishing the whole cell configuration. Cell attached recordings were carried out in the same recording solution in the absence of any blockers unless otherwise stated at zero current injection.

IPSCs and action potentials were analysed using WinEDR (ver 4.0.2) and WinWCP (ver 5.7.0) and frequency and amplitude of IPSCs were measured from 60 s epochs after allowing enough time for internal solution to diffuse into the cells. Tonic currents were measured as the difference in stable membrane current from 10 s epochs before and after the application of 59 μM bicuculline. For IPSC kinetics, T_50_, weighted decay time, rise time and charge transfer were measured from single uncontaminated IPSCs^80^.

Weighted decay times have been reported by factoring in mono- and bi-exponentially decaying events:

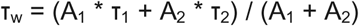

where τ_1_ and τ_2_ are exponential decay time constants, and A_1_ and A_2_ are the relative amplitude contributions of τ_1_ and τ_2_, respectively.

### Immunofluorescence staining

Adult (5–6 months of age) mice were sacrificed by a lethal dose of sodium pentobarbital delivered intraperitoneally, then quickly perfused with 4% paraformaldehyde (PFA) transcardially followed by brain removal. Brains were post-fixed in PFA for a further 2-4 hr at 4°C followed by transfer to a cryoprotectant solution comprising: 0.1 M phosphate buffer, 0.03% w/ v sodium azide and 30% w/ v sucrose and incubation at 4°C until the tissue sank. Coronal sections (40 µm) were cut from frozen brain using a Leica SM200R sliding microtome (Leica Microsytems, Germany). Brain sections were collected in 0.1 M phosphate buffer with 0.03% w/v sodium azide and were stored at 4°C for further processing.

Groups of every 12^th^ slice of the hippocampus were washed with phosphate buffered saline (PBS) and incubated for 1 hr at room temperature in permeabilization solution containing 0.5% bovine serum albumin, 2% normal goat serum, 0.2% triton-x-100 in PBS. Brain sections were incubated with primary antibodies (1: 1000 dilution; GABA, Millipore – ABN131; parvalbumin, Abcam – ab11427) for 2 hr at 4°C in the same permeabilization solution followed by three washes in PBS at room temperature for 15 min with gentle shaking. Secondary antibody (ThermoFisher, A-11008) was applied to slices for 3 hr at room temperature. After four washes in PBS sections were mounted on microscope slides (VWR International) in ProLong Gold Antifade reagent (ThermoFisher).

Brains sections were imaged with a Leica SP8 or Zeiss Axioscop LSM510 confocal microscope using a x40 oil objective. Images were captured as tiles and z-stacks and analysed using Image J (Ver 1.54f) by counting the number of immunofluorescence positive somata within the specific areas of the hippocampus.

### 3D reconstruction of cells and Sholl analysis

During some of our acute brain slice recordings, we included lucifer yellow (0.5 mg/ ml; ThermoFisher, L453) in our internal solution allowing the neural projections and soma to be filled with dye during our recordings. Upon completion of recordings the patch pipette was gently raised above the cell to allow the cells to re-seal and slices were quickly fixed by transferring to 4% PFA at 4°C overnight. Brain slices were washed 4 times with PBS and mounted in glass microscope slides in ProLong Gold. Dye-filled neurons were identified under the microscope and imaged using a Zeiss Axioscop LSM510 confocal microscope using a x25 oil objective as z-stacks in 3D. Images were segmented and Sholl analysis in 3D was carried out in Image J (ver 1,54f)

### Peak-scaled Non-Stationary Noise Analysis (PS-NSNA)

PS-NSNA was performed on the IPSC decay from the peak to the end of the IPSC decay. The average current was scaled to the peak of the individual IPSC amplitudes, and the amplitude was divided into bins of equal size. Fluctuations in current resulting from channel opening and closing were identified by subtracting the mean current variance from the mean IPSC current decay. The amplitude variance within each scaled average bin was then calculated and plotted against the average bin current^80,81^. The variance - mean current amplitude relationship was fitted by:

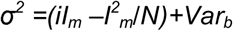

where σ^2^ is the current variance, *i* represents the unitary current, *I_m_* is mean current and *N* is the average number of receptor channels open in response to a single vesicle release in the synapse. *Var_b_* is the baseline variance. This plot estimated single channel current (pA) by the initial gradient of the parabola, which was converted to unitary conductance by

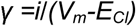

where γ is the unitary conductance, *V_m_*is the holding potential of the cell, *E_Cl_* is the Cl^-^ reversal potential and *i* represents the single channel current.

### Experimental design and statistics

All statistical tests that have been used, and applied to sample sizes in the study, are indicated in the figure legends and in results. For comparing two samples, normally distributed data and nonparametric data, we used a two-tailed unpaired t-test and Mann-Whitney test, respectively. Three or more groups were compared using a one-way ANOVA (normally distributed) or Kruskal-Wallis (KW) ANOVA (nonparametric distribution) followed by a Tukey or Dunn’s multiple comparisons test respectively. All statistical tests were carried out (including test for normality) using GraphPad Instat 3 and SPSS (ver 24). Graphs were plotted in Origin 2021b. Data in the bar charts represent mean ± standard error of mean. Data in box plots show 25%-75% interquartile ranges, 5-95% whiskers and the median.

## Supporting information

Supplementary details

## Acknowledgements

TGS, EMCF and VLJT were funded by a Wellcome Trust Collaborative Award in Science (217199/Z/19/Z). VLJT was supported by the Francis Crick Institute which receives its core funding from Cancer Research UK (CC2080), the UK Medical Research Council (CC2080), and the Wellcome Trust (CC2080). SH was a recipient of an International Rett Syndrome Foundation Fellowship (3606).

## Author contributions

Conceptualization – SBH, EMCF, VLJT, TGS. Data acquisition – SH. Generation of mouse lines – ELE, SWS. Project leadership and funding acquisition - SBH, EMCF, VLJT, TGS. Initial draft of the manuscript was written by SBH, VLJT, TGS and all authors contributed to the writing of the manuscript.

## Materials & Correspondence

All data are available in the main manuscript or supplementary materials. Additional requests for data and materials in the manuscript will be made available upon reasonable request by the corresponding authors under suitable materials transfer agreements (MTAs).

---

**Supplementary figure 1.**
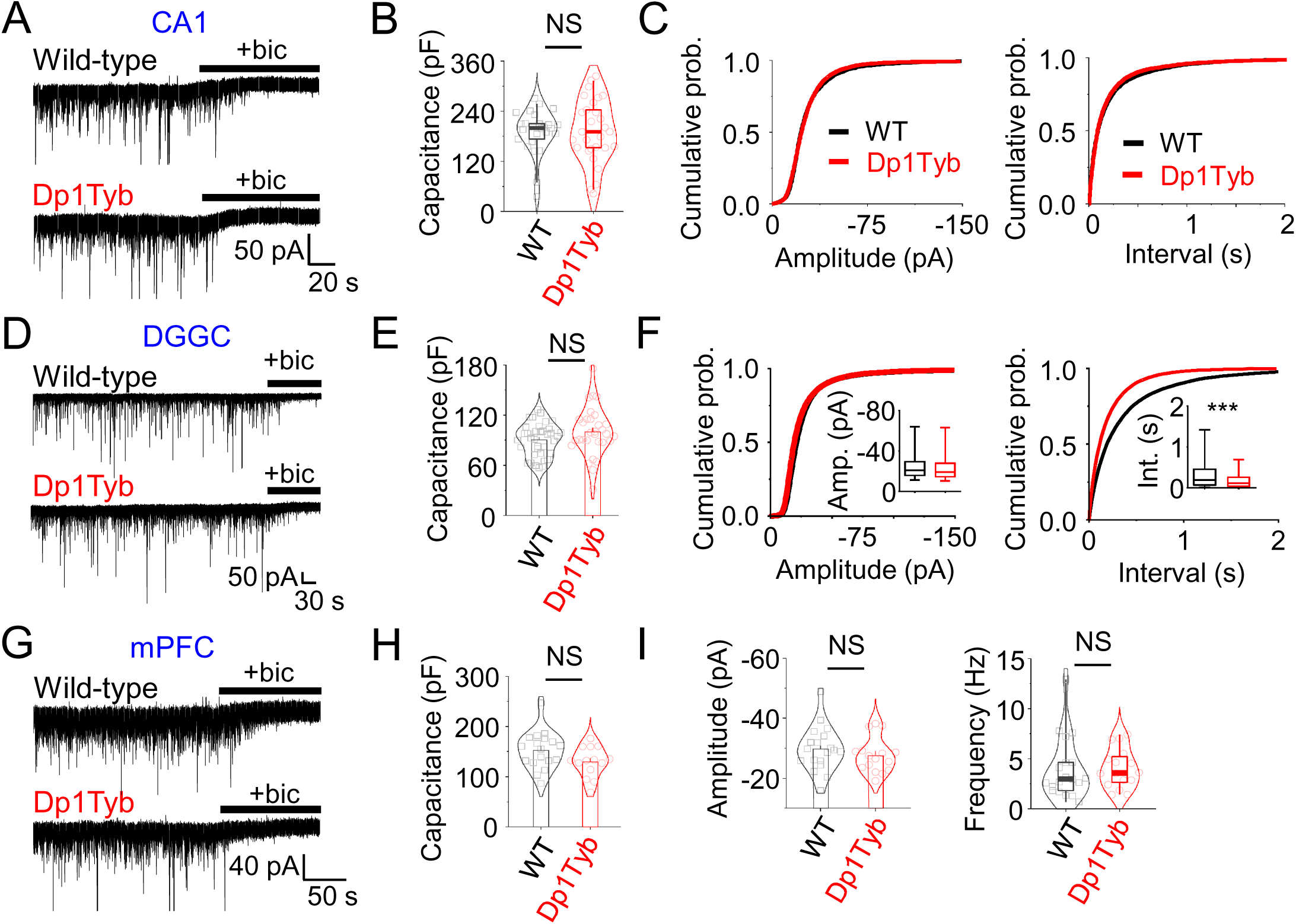
Phasic inhibition in the Dp1Tyb mouse model of Down syndrome. *A,* Representative spontaneous inhibitory current (sIPSC) recordings of CA1 pyramidal neurons from wild-type and Dp1Tyb mice. Addition of bicuculline (50 µM; +bic) blocks postsynaptic currents confirming their GABAergic identity. *B,* Whole cell capacitance of CA1 pyramidal neurons. *C,* Cumulative probability distributions of sIPSC amplitudes and interevent intervals of CA1 neurons. *D,* Representative sIPSC recordings of dentate gyrus granule cells (DGGCs) from wild-type and Dp1Tyb mice. *E,* Whole cell capacitance of DGGCs. *F,* Cumulative probability distributions of sIPSC amplitude and interevent intervals of DGGCs. Insets, box-plots showing unchanged median amplitude but lower inter event interaval (int.) for Dp1Tyb sIPSCs compared to wild-type. *G,* Representative sIPSC recordings of layer II/ III medial prefrontal cortex (mPFC) neurons from wild-type and Dp1Tyb mice. *H,* Whole cell capacitance of mPFC neurons. *I,* There was no change in sIPSC amplitudes and frequency of mPFC neurons when averages or medians per cell was analysed although their distributions were marginally affected (see main figures). NS – not significant, ***P<0.001, two-tailed unpaired t-test or Mann-Whitney test. n = 14-36 cells; 2205-3465 events for distributions, 7-9 animals.

**Supplementary figure 2.**
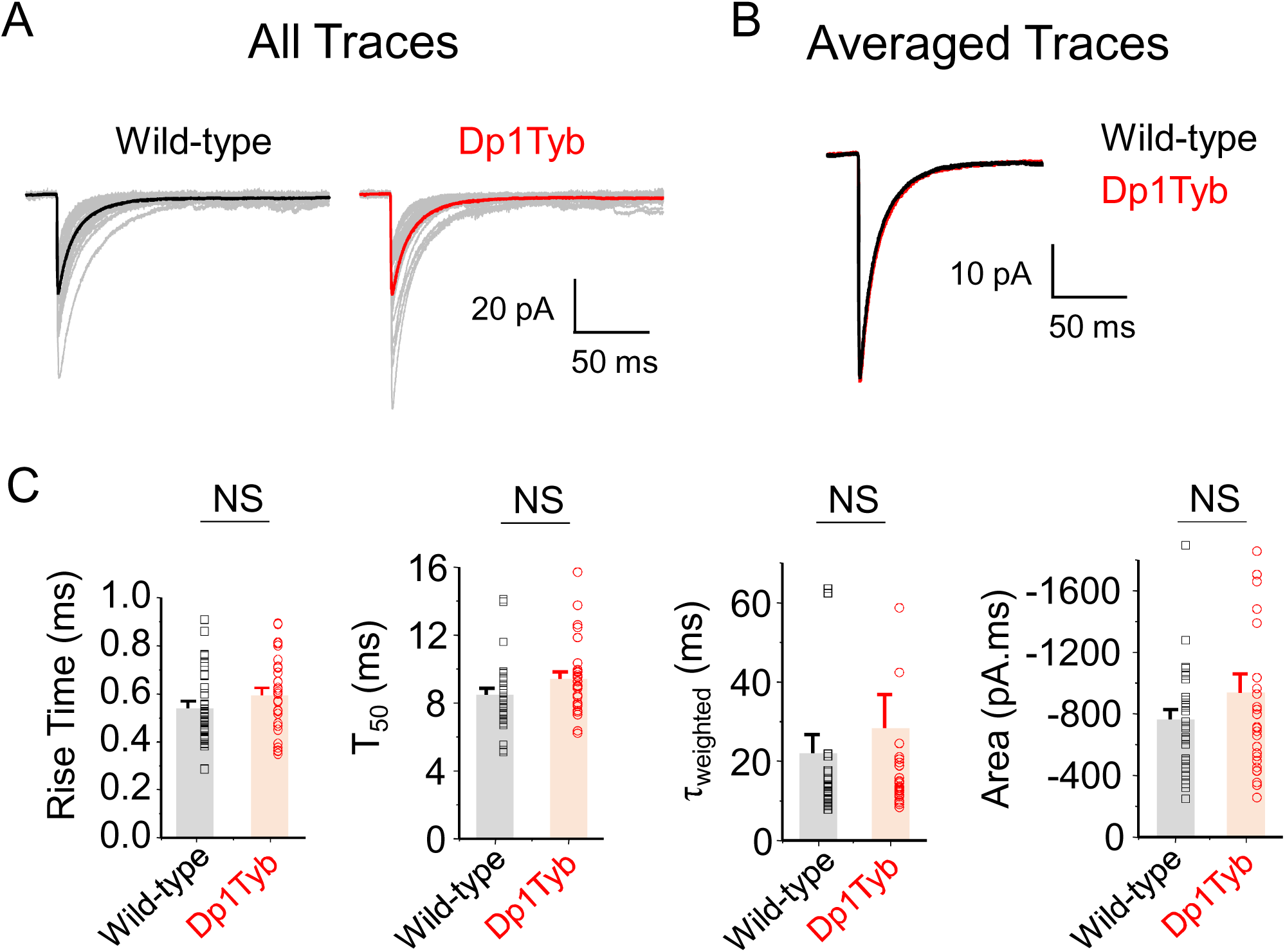
Unchanged IPSC kinetics in Dp1Tyb dentate granule neurons. *A,* sIPSC waveforms of wild-type and Dp1Tyb dentate gyrus granule neurons. Averaged waveforms have been depicted in darker shades. *B,* Overlay of average wild-type and Dp1Tyb sIPSC waveforms. C, Average rise-times. T_50_, weighted τ and charge transfer of IPSC waveforms. NS – not significant, two-tailed unpaired t-test. n = 29-31 cells; 8-9 animals

**Supplementary figure 3.**
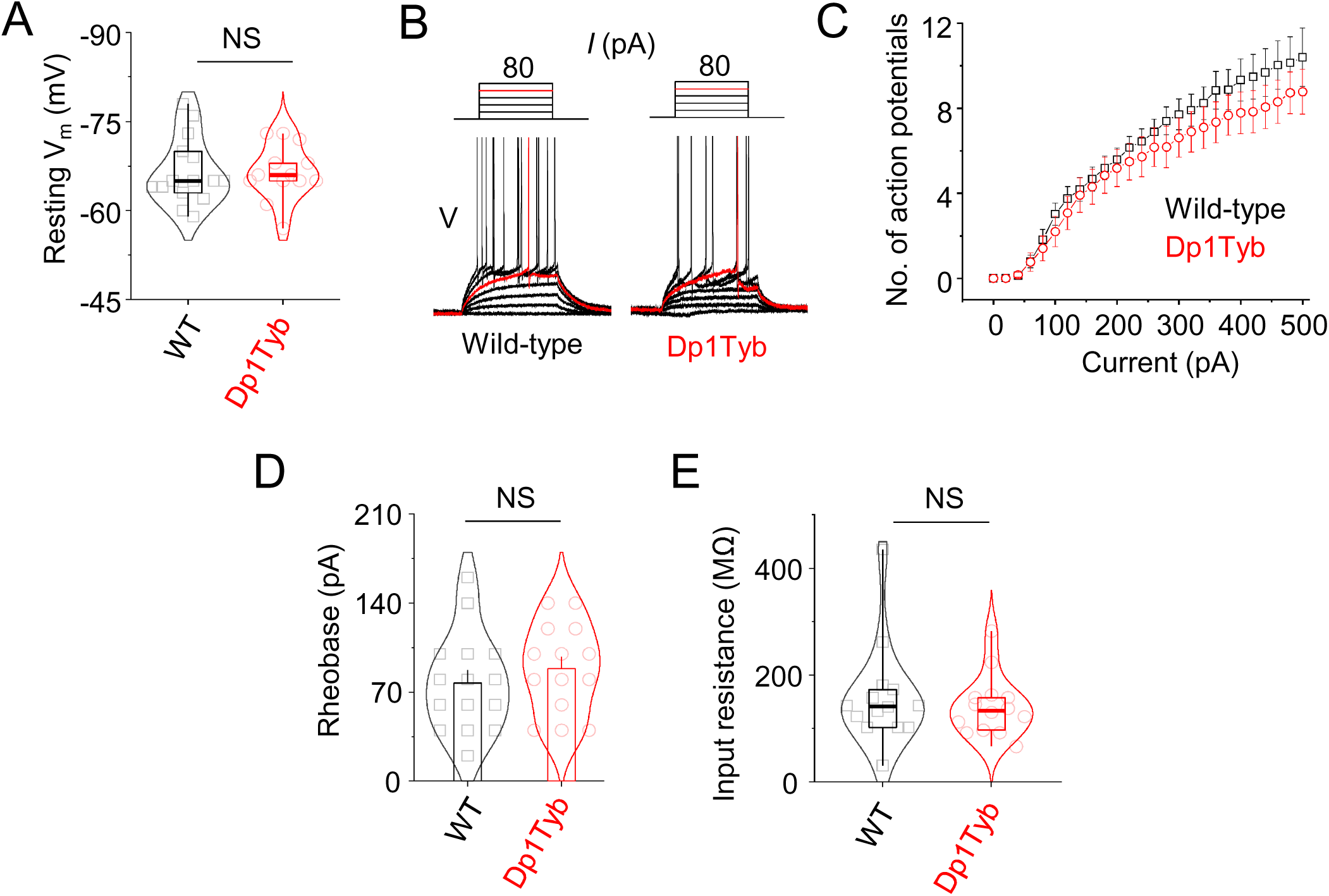
Unchanged spike firing properties in Dp1Tyb CA1 neurons. *A,* Resting membrane potentials of wild-type (WT) and Dp1Tyb CA1 neurons. *B,* Example current clamp recordings of wild-type and Dp1Tyb CA1 neurons in response to step current injection. The rheobase trace has been depicted in red. *C,* Current-action potential relationship of wild-type and Dp1Tyb CA1 neurons. *D,* Rheobase of wild-type and Dp1Tyb CA1 neurons. *E,* Input resistance of wild-type and Dp1Tyb CA1 neurons. NS – not significant, two-tailed unpaired t-test or Mann-Whitney test. n = 13-15 cells; 3-4 animals.

**Supplementary figure 4.**
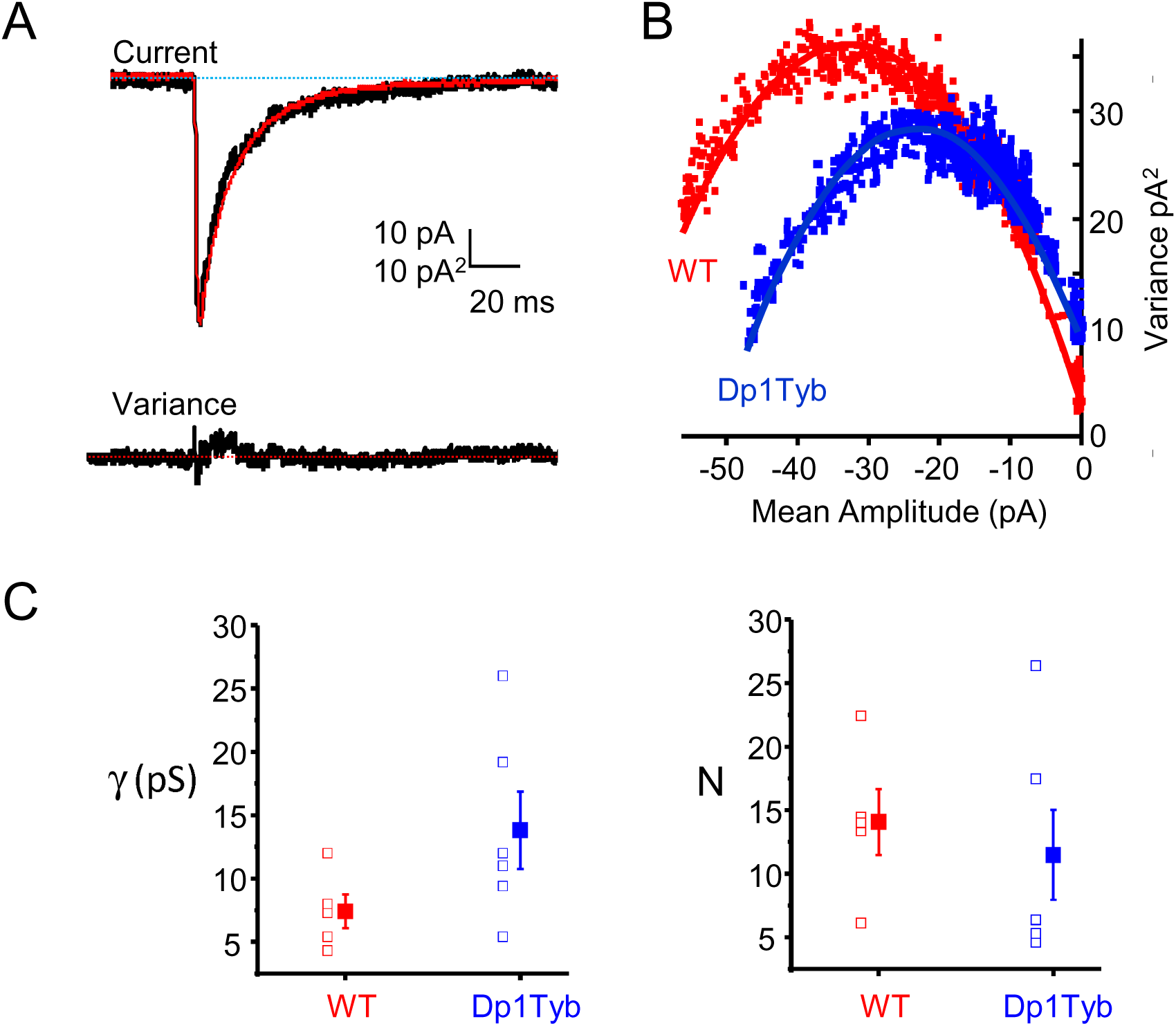
Synaptic current noise analysis in DGGC. *A,* Representative voltage clamp IPSC recording (black) compared to the mean IPSC (red; upper trace) and associated variance (lower trace) from a wild-type dentate gyrus granule cell (DGGC). *B*, Mean current variance relationships for wild-type and Dp1Tyb representative DGGCs. *C,* Estimates for single channel conductance (γ) and the number of synaptic receptors (N) obtained from PS-NSNA for IPSCs from wild-type and Dp1Tyb DGGCs (n = 5 - 6). Filled symbols represent mean +/- sem. Unpaired t-test reveals P>0.05 and P >0.05 for channel conductance and receptor number comparison between wild-type and Dp1Tyb cells.

**Supplementary figure 5.**
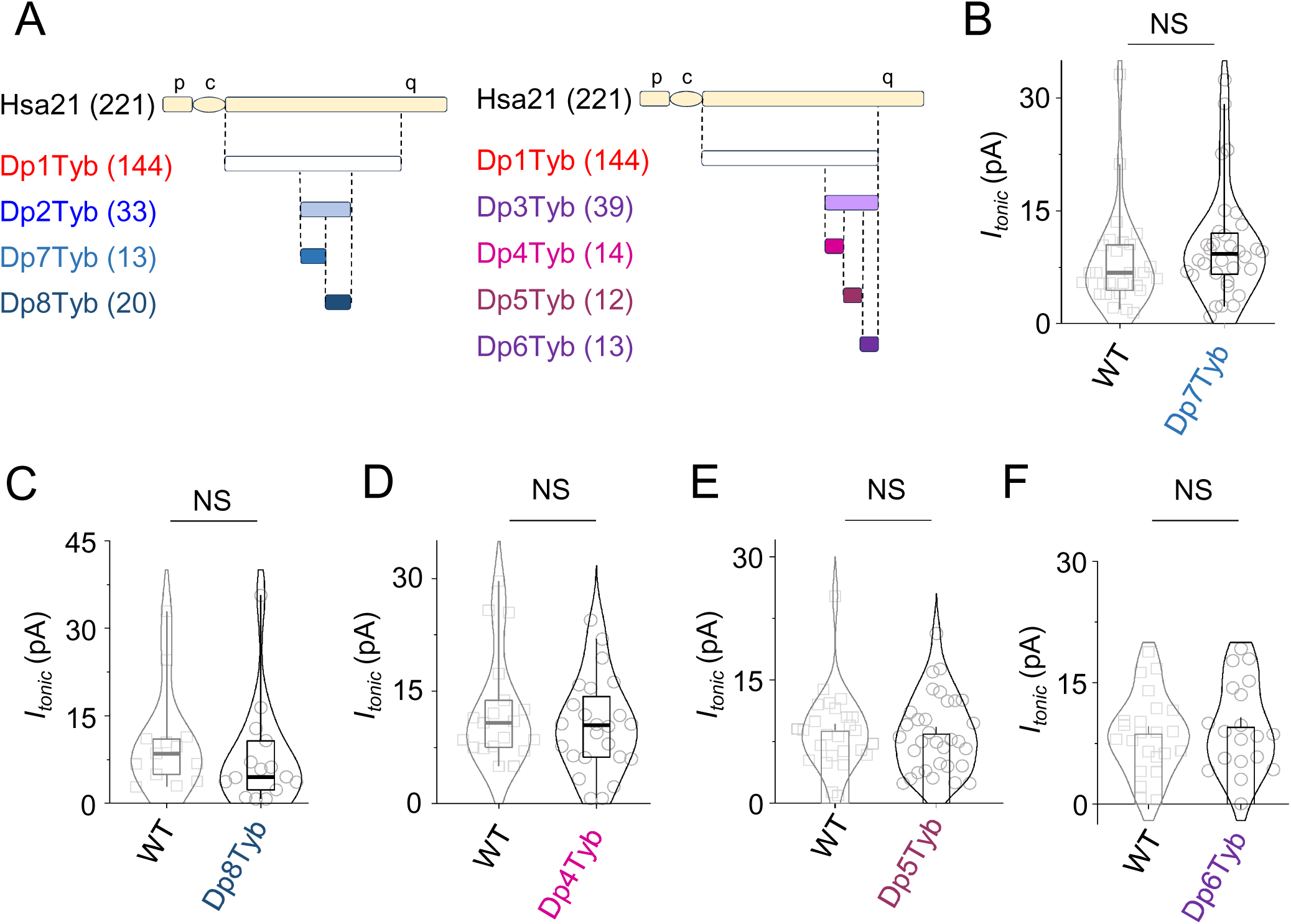
Breaking up the Dp2Tyb or Dp3Tyb regions causes the tonic inhibition phenotype to disappear. *A,* Schematic showing the short (p) and long (q) arms and the centromere (c) of Hsa21 aligned with orthologous regions on Mmu16 that are present in a third copy in Dp1Tyb, Dp2Tyb, Dp3Tyb, Dp4Tyb, Dp5Tyb, Dp6Tyb, Dp7Tyb and Dp8Tyb mice. Numbers in parentheses indicate numbers of coding genes in these intervals. *B-F,* Tonic currents of wild-type (WT) and Dp7Tyb (B), Dp8Tyb (C), Dp4Tyb (D), Dp5Tyb (E) or Dp6Tyb (F) dentate gyrus granule neurons. NS – not significant, two-tailed unpaired t-test or Mann-Whitney test. n = 13-32 cells; 5-13 animals.

