## Supplementary details for "Hippocampal circuit-specific enhancement of GABA-inhibition caused by discrete gene regions in a Down syndrome model"

Supplementary Table 1 - Dp mouse models and gene counts

| Gene ID | Gene name | Gene start (bp) | Gene end (bp) | | | | |
|---|---|---|---|---|---|---|---|
| ENSMUSG00000032948 | Lipi | 75337402 | 75382949 | | | | |
| ENSMUSG00000032940 | Rbm11 | 75389732 | 75399717 | | | | |
| ENSMUSG00000032932 | Hspa13 | 75542319 | 75563992 | | | | |
| ENSMUSG00000022876 | Samsn1 | 75655681 | 75819169 | | | | |
| ENSMUSG00000048490 | Nrip1 | 76084288 | 76170715 | | | | |
| ENSMUSG00000022867 | Usp25 | 76810594 | 76913668 | | | | |
| ENSMUSG00000022865 | Cxadr | 78098377 | 78156662 | | | | |
| ENSMUSG00000022863 | Btg3 | 78129525 | 78174080 | | | | |
| ENSMUSG00000022864 | D16Ertd472e | 78337226 | 78373596 | | | | |
| ENSMUSG00000022860 | Chodl | 78727836 | 78748621 | | | | |
| ENSMUSG00000022857 | Tmprss15 | 78749896 | 78887985 | | | | |
| ENSMUSG00000022762 | Ncam2 | 80997585 | 81423716 | | | | |
| ENSMUSG00000022889 | Mrpl39 | 84514464 | 84532630 | | | | |
| ENSMUSG00000053062 | Jam2 | 84571011 | 84622816 | | | | |
| ENSMUSG00000022890 | Atp5pf | 84624754 | 84632513 | | | | |
| ENSMUSG00000008976 | Gabpa | 84631813 | 84660667 | | | | |
| ENSMUSG00000022892 | App | 84746573 | 84970654 | | | | |
| ENSMUSG00000041134 | Cyyr1 | 85218421 | 85350285 | | | | |
| ENSMUSG00000022893 | Adamts1 | 85590715 | 85600001 | | | | |
| ENSMUSG00000022894 | Adamts5 | 85653061 | 85698716 | | | | |
| ENSMUSG00000044442 | N6amt1 | 87151073 | 87165630 | | | | |
| ENSMUSG00000052299 | Ltn1 | 87173539 | 87229500 | | | | |
| ENSMUSG00000041079 | Rwdd2b | 87230295 | 87237461 | | | | |
| ENSMUSG00000025616 | Usp16 | 87251591 | 87280405 | | | | |
| ENSMUSG00000025613 | Cct8 | 87280214 | 87292761 | | | | |
| ENSMUSG00000025610 | Map3k7cl | 87350218 | 87392224 | | | | |
| ENSMUSG00000025612 | Bach1 | 87495833 | 87530234 | | | | |
| ENSMUSG00000022935 | Grik1 | 87692788 | 88087153 | | | | |
| ENSMUSG00000055811 | Cldn17 | 88302695 | 88303866 | | | | |
| ENSMUSG00000050520 | Cldn8 | 88357716 | 88360071 | | | | |
| ENSMUSG00000050239 | Krtap24-1 | 88407593 | 88409167 | | | | |
| ENSMUSG00000045331 | 2310079G19Rik | 88423674 | 88424554 | | | | |
| ENSMUSG00000071471 | Krtap26-1 | 88443712 | 88444684 | | | | |
| ENSMUSG00000090515 | Krtap27-1 | 88467914 | 88468559 | | | | |
| ENSMUSG00000050704 | Krtap13-22 | 88504057 | 88504850 | | | | |
| ENSMUSG00000056350 | Krtap13-1 | 88525750 | 88526499 | | **Dp9Tyb (72 genes)** | | |
| ENSMUSG00000050224 | Krtap13 | 88547632 | 88548523 | | | | |
| ENSMUSG00000068078 | Krtap13-20 | 88555735 | 88556680 | | | | |
| ENSMUSG00000048830 | Krtap13-21 | 88570067 | 88571094 | | | | |
| ENSMUSG00000009047 | Krtap13-23 | 88574784 | 88575616 | | | | |
| ENSMUSG00000116636 | Krtap13-24 | 88608328 | 88609108 | | | | |
| ENSMUSG00000022931 | Krtap15-1 | 88617103 | 88626732 | | | | |
| ENSMUSG00000074928 | Krtap14 | 88622178 | 88623033 | | | | |
| ENSMUSG00000060691 | Krtap19-1 | 88665805 | 88666299 | | | | |
| ENSMUSG00000057650 | Krtap19-2 | 88670515 | 88671157 | | | | |
| ENSMUSG00000060469 | Krtap19-3 | 88674401 | 88674926 | | | | |
| ENSMUSG00000043982 | Krtap19-4 | 88681674 | 88681977 | | | | |
| ENSMUSG00000051802 | Krtap19-5 | 88692856 | 88693345 | | | | |
| ENSMUSG00000057174 | Krtap19-9b | 88728692 | 88729153 | | | | |
| ENSMUSG00000096534 | Krtap16-3 | 88759161 | 88759761 | | | | |
| ENSMUSG00000095992 | Krtap22-2 | 88807283 | 88807647 | | | | |
| ENSMUSG00000068075 | Krtap6-3 | 88812172 | 88812734 | | | | |
| ENSMUSG00000058172 | Krtap6-1 | 88828587 | 88829180 | | | | |
| ENSMUSG00000068074 | Krtap6-7 | 88837811 | 88838360 | | | | |
| ENSMUSG00000062400 | Krtap6-5 | 88844177 | 88844771 | | | | |
| ENSMUSG00000068073 | Krtap6-6 | 88860300 | 88860890 | | | | |
| ENSMUSG00000057855 | Krtap20-1 | 88880903 | 88881422 | | | | |
| ENSMUSG00000068071 | krtap20-23 | 88937739 | 88938227 | | | | |
| ENSMUSG00000059113 | Krtap20-20 | 88947974 | 88948438 | | | | |
| ENSMUSG00000044227 | Krtap20-21 | 88954840 | 88955236 | | | | |
| ENSMUSG00000068068 | Krtap20-22 | 88966378 | 88966536 | | | | |
| ENSMUSG00000068067 | Krtap20-24 | 88971623 | 88972087 | | | | |
| ENSMUSG00000091039 | Krtap20-2 | 89002749 | 89003282 | | | | |
| ENSMUSG00000058368 | Krtap21-1 | 89200106 | 89200756 | | | | |
| ENSMUSG00000062433 | Krtap6-2 | 89216214 | 89216999 | | | | |
| ENSMUSG00000059632 | Krtap8-1 | 89284267 | 89284840 | | | | |
| ENSMUSG00000056706 | Krtap7-1 | 89304591 | 89305211 | | | | |
| ENSMUSG00000091212 | Krtap11-1 | 89367064 | 89368071 | | | | |
| ENSMUSG00000002489 | Tiam1 | 89583999 | 89940657 | | | | |
| ENSMUSG00000022982 | Sod1 | 90017642 | 90023217 | | | | |
| ENSMUSG00000022983 | Scaf4 | 90022568 | 90081391 | | | | |
| ENSMUSG00000053414 | Hunk | 90182901 | 90296441 | **Dp1Tyb (144 genes)** | | | |
| ENSMUSG00000022978 | Mis18a | 90516200 | 90524292 | | | | |
| ENSMUSG00000039956 | Mrap | 90535095 | 90546673 | | | | |
| ENSMUSG00000039929 | Urb1 | 90548415 | 90607301 | | | | |
| ENSMUSG00000039903 | Eva1c | 90623607 | 90701997 | | | | |
| ENSMUSG00000022972 | Cfap298 | 90722697 | 90732002 | | | | |
| ENSMUSG00000022973 | Synj1 | 90732980 | 90808196 | | | | **Ts65Dn (132 genes)** |
| ENSMUSG00000022974 | Paxbp1 | 90810925 | 90841431 | | | **Dp7Tyb (13 genes)** | |
| ENSMUSG00000039851 | Epcip | 90850823 | 90892010 | | | | |
| **ENSMUSG00000039830** | **Olig2** | **91022345** | **91025565** | | | | |
| **ENSMUSG00000046160** | **Olig1** | **91066660** | **91068821** | | | | |
| ENSMUSG00000022971 | Ifnar2 | 91169671 | 91202477 | | | | |
| ENSMUSG00000093701 | Gm21970 | 91180749 | 91222722 | | | | |
| ENSMUSG00000022969 | Il10rb | 91203052 | 91222722 | | | | |
| ENSMUSG00000022967 | Ifnar1 | 91282126 | 91304329 | | | | |
| ENSMUSG00000022965 | Ifngr2 | 91343960 | 91362511 | | | | |
| ENSMUSG00000022964 | Tmem50b | 91371391 | 91394688 | | | | |
| ENSMUSG00000039763 | Dnajc28 | 91411142 | 91415914 | | **Dp2Tyb (33 genes)** | | |
| ENSMUSG00000022962 | Gart | 91418074 | 91443840 | | | | |
| ENSMUSG00000022961 | Son | 91444394 | 91476109 | | | | |
| ENSMUSG00000022960 | Donson | 91473696 | 91485658 | | | | |
| ENSMUSG00000058240 | Cryzl1 | 91486210 | 91525863 | | | | |
| ENSMUSG00000022957 | Itsn1 | 91526169 | 91717485 | | | | |
| ENSMUSG00000022956 | Atp5po | 91722102 | 91728575 | | | **Dp8Tyb (20 genes)** | |
| ENSMUSG00000039680 | Mrps6 | 91855158 | 91909115 | | | | |
| ENSMUSG00000089774 | Slc5a3 | 91855210 | 91884361 | | | | |
| ENSMUSG00000039672 | Kcne2 | 92089277 | 92095017 | | | | |
| ENSMUSG00000051989 | Smim11 | 92098174 | 92109929 | | | | |
| ENSMUSG00000051728 | Fam243 | 92115653 | 92118329 | | | | |
| ENSMUSG00000116967 | Smim34 | 92119046 | 92142068 | | | | |
| ENSMUSG00000039639 | Kcne1 | 92142870 | 92156356 | | | | |
| ENSMUSG00000022951 | Rcan1 | 92188841 | 92267755 | | | | |
| ENSMUSG00000022949 | Clic6 | 92282624 | 92338131 | | | | |
| ENSMUSG00000022952 | Runx1 | 92398354 | 92623037 | | | | |
| ENSMUSG00000022948 | Setd4 | 93380345 | 93400951 | | | | |
| ENSMUSG00000051483 | Cbr1 | 93402741 | 93407393 | | | | |
| ENSMUSG00000082815 | Cbr1b | 93424985 | 93427339 | | | | |
| ENSMUSG00000022947 | Cbr3 | 93480103 | 93487878 | | | | |
| ENSMUSG00000022946 | Dop1b | 93508792 | 93607478 | | | | |
| ENSMUSG00000039456 | Morc3 | 93629009 | 93672961 | | | | |
| ENSMUSG00000022945 | Chaf1b | 93680789 | 93703003 | | | **Dp4Tyb (14 genes)** | |
| ENSMUSG00000047109 | Cldn14 | 93715919 | 93809696 | | | | |
| ENSMUSG00000062713 | Sim2 | 93885790 | 93927891 | | | | |
| ENSMUSG00000040820 | Hlcs | 93929741 | 94114430 | | | | |
| ENSMUSG00000022941 | Ripply3 | 94129279 | 94137794 | | | | |
| ENSMUSG00000022940 | Pigp | 94159622 | 94172701 | | | | |
| ENSMUSG00000040785 | Ttc3 | 94171477 | 94270202 | | | | |
| ENSMUSG00000022898 | Vps26c | 94298642 | 94327689 | | | | |
| **ENSMUSG00000022897** | **Dyrk1a** | **94370869** | **94496376** | | | | |
| ENSMUSG00000043301 | Kcnj6 | 94549495 | 94798560 | | | | |
| ENSMUSG00000062609 | Kcnj15 | 95058417 | 95101119 | | | | |
| ENSMUSG00000040732 | Erg | 95160028 | 95387452 | | | | |
| ENSMUSG00000022895 | Ets2 | 95502942 | 95522095 | | | | |
| ENSMUSG00000022913 | Psmg1 | 95781133 | 95792160 | | **Dp3Tyb (39 genes)** | **Dp5Tyb (12 genes)** | |
| ENSMUSG00000022914 | Brwd1 | 95793292 | 95883726 | | | | |
| ENSMUSG00000040681 | Hmgn1 | 95921818 | 95928929 | | | | |
| ENSMUSG00000023147 | Get1 | 95946607 | 95959052 | | | | |
| ENSMUSG00000045275 | Lca5l | 95959607 | 95993471 | | | | |
| ENSMUSG00000040666 | Sh3bgr | 96001650 | 96030135 | | | | |
| ENSMUSG00000074892 | B3galt5 | 96037001 | 96121059 | | | | |
| ENSMUSG00000000159 | Igsf5 | 96162868 | 96223321 | | | | |
| ENSMUSG00000000157 | Itgb2l | 96223488 | 96244819 | | | | |
| ENSMUSG00000090223 | Pcp4 | 96268806 | 96326993 | | | | |
| ENSMUSG00000050272 | Dscam | 96392040 | 96971952 | | | | |
| ENSMUSG00000040605 | Bace2 | 97157942 | 97244136 | | | | |
| ENSMUSG00000000386 | Mx1 | 97248235 | 97264107 | | | | |
| ENSMUSG00000022938 | Fam3b | 97272165 | 97316016 | | | **Dp6Tyb (13 genes)** | |
| ENSMUSG00000023341 | Mx2 | 97336508 | 97362100 | | | | |
| ENSMUSG00000000385 | Tmprss2 | 97365882 | 97412395 | | | | |
| ENSMUSG00000005251 | Ripk4 | 97543133 | 97564987 | | | | |
| ENSMUSG00000014039 | Prdm15 | 97592667 | 97653050 | | | | |
| ENSMUSG00000045975 | C2cd2 | 97656409 | 97763798 | | | | |
| ENSMUSG00000046962 | Zbtb21 | 97744557 | 97763822 | | | | |

**Protein coding genes in the Dp mouse strains**

Protein coding genes downloaded using Biomart function of Ensembl on mouse genome assembly GRCm39
List shows Mmu16 genes present in three copies in each of the Dp strains, gving gene ID, name and start and end coordinates
Extent of region present in three copies in each strain is indicated on the right.
Numbers of Mmu16 genes present in three copies in each of the Dp strains and in Ts65Dn mice is given in parentheses for each strain.
Ts65Dn mice have a further 46 genes on Mmu17 that are in 3 copies that are not shown.
The three candidate genes tested (Olig1, Olig2 and Dyrk1a) are indicated in bold.
Date of download: 19/02/2025
