## supplementary data for "Hippocampal circuit-specific enhancement of GABA-inhibition caused by discrete gene regions in a Down syndrome model"

**Supplementary Table 2 – Phasic and tonic inhibition in Down syndrome mouse lines**

| | Freq (Hz) | | P Value | Amp (pA) | | P Value | Tonic (pA) | | P Value | WCell Cap | | P Value |
|---|---|---|---|---|---|---|---|---|---|---|---|---|
| | Wild-type | Dp | | Wild-type | Dp | | Wild-type | Dp | | Wild-type | Dp | |
| CA1 Dp1Tyb | 7.2 ± 1.3 (25) | 8.9 ± 1.1 (30) | 0.2205 | -28.6 ± 1 (25) | -27.7 ± 1.4 (30) | 0.6398 | 19.6 ± 2.2 (21) | 20.2 ± 2.1 (24) | 0.9728 | 191 ± 10.6 (25) | 192.4 ± 14 (30) | 0.9447 |
| mPFC Dp1Tyb | 3.9 ± 3 (19) IEI: 0.177 s (3117) | 4 ± 0.5 (16) IEI: 0.215 s (2222) | 0.4765 | -29.7 ± 1.7 (19) Median: -25.3 (3465) | -27.5 ± 1.4 (16) Median: -23.3 (2205) | 0.3384 | 11.2 ± 2.1 (16) | 10.9 ± 2 (13) | 0.9227 | 152.8 ± 8.7 (18) | 129.5 ± 8 (14) | 0.0651 |
| DG Dp1Tyb | 3.4 ± 0.3 (29) | 4.7 ± 0.5 (31) | 0.0413 | -26.8 ± 1 (29) | -25.2 ± 1.2 (31) | 0.1684 | 6.1 ± 0.8 (36) | 9.6 ± 1 (37) | 0.0180 | 90.5 ± 3.2 (36) | 100 ± 5.1 (31) | 0.1088 |
| DG Dp2Tyb | 2.2 ± 0.4 (18) | 3.4 ± 0.3 (29) | 0.002 | -29 ± 1.9 (18) | -25.1 ± 1.1 (29) | 0.0614 | 6.3 ± 1 (15) | 9.3 ± 1 (27) | 0.0489 | 85.8 ± 5 (18) | 88.1 ± 5 (31) | 0.7671 |
| DG Dp3Tyb | 2.6 ± 0.3 (37) | 4 ± 0.4 (36) | 0.0141 | -23.4 ± 1.2 (37) | -23.8 ± 1.4 (36) | 0.9165 | 7.4 ± 0.8 (30) | 10.8 ± 1 (28) | 0.0117 | 88.3 ± 3.5 (37) | 97.4 ± 3.4 (40) | 0.0673 |
| DG Dp9Tyb | 2.1 ± 0.2 (20) | 2.5 ± 0.5 (16) | 0.8861 | -29.4 ± 1.7 (20) | -29.3 ± 1.7 (16) | 0.7623 | 6.3 ± 0.8 (18) | 8.1 ± 1.8 (14) | 0.3162 | 90 ± 4.2 (20) | 93 ± 7.4 (17) | 0.9878 |
| DG Dp4Tyb | 2.9 ± 0.4 (23) | 3.1 ± 0.4 (30) | 0.5842 | -34.9 ± 1.8 (23) | -31.2 ± 1.3 (30) | 0.0834 | 12.4 ± 1.6 (19) | 10.4 ± 1.3 (24) | 0.4411 | nd | | |
| DG Dp5Tyb | 2.6 ± 0.4 (32) | 2.7 ± 0.3 (37) | 0.4593 | -29.9 ± 1.7 (32) | -28 ± 1.1 (37) | 0.5043 | 8.7 ± 0.9 (26) | 8.4 ± 0.8 (32) | 0.8126 | | | |
| DG Dp6Tyb | 2.4 ± 0.3 (26) | 2.3 ± 0.2 (24) | 0.6682 | -31.4 ± 1.5 (26) | -29.8 ± 1.5 (24) | 0.4915 | 8.6 ± 1 (23) | 9.5 ± 1.3 (19) | 0.6037 | | | |
| DG Dp7Tyb | 3 ± 0.4 (33) | 2.5 ± 0.3 (35) | 0.5315 | -28.5 ± 1.1 (33) | -27.9 ± 1.3 (35) | 0.4180 | 7.2 ± 0.8 (26) | 10.5 ± 1.3 (32) | 0.0535 | | | |
| DG Dp8Tyb | 3.8 ± 0.6 (15) | 3.2 ± 0.4 (22) | 0.2589 | -32.3 ± 2.4 (15) | -34 ± 2 (22) | 0.5953 | 10.5 ± 2.4 (13) | 7.7 ± 2.3 (15) | 0.1846 | | | |

Frequency (Freq) and amplitude (Amp) of phasic inhibition along with tonic currents and whole-cell capacitance (WCell Cap) of duplication (Dp) mouse models compared to littermate wild-type controls. Number in brackets show n numbers of cells or events. P values are from two-tailed unpaired t-tests or Mann-Whitney test and can be found in figures and their legends. cornu amonis area 1, CA1; dentate gyrus, DG; medial prefrontal cortex, mPFC; not determined, nd

**Supplementary Table 3 – Current clamp properties of Down syndrome mouse lines**

| | Dentate gyrus granule cells | | CA1 pyramidal neurons | |
|---|---|---|---|---|
| | Wild-type | Dp1Tyb | Wild-type | Dp1Tyb |
| Vm (mV) | -75.3 ± 1.2 (28) | -77.5 ± 0.8 (26) | -66.5 ± 1.4 (15) | -66.5 ± 1.3 (13) |
| Rheobase (pA) | 68.6 ± 3.8 (28) [in bicuculline: 66.1 ± 4.7 (13)] | 88 ± 4.8 (26) [in bicuculline: 61.7 ± 6.2 (12)] | 77.3 ± 9.9 (15) | 88.6 ± 9.3 (14) |
| Input Resistance (MΩ) | 500 ± 360 (28) [in bicuculline: 589 ± 757 (11)] | 359 ± 272 (26) [in bicuculline: 482 ± 512 (11)] | 140 ± 263 (14) | 141 ± 151 (14) |

Active and passive membrane properties of dentate gyrus granule cells and CA1 pyramidal neurons. Vm, resting membrane potential.

**Supplementary Table 4 - Phasic and tonic inhibition in gene-copy rescued Down syndrome dentate granule cells**

| | | WT | Dp2Tyb or Dp3Tyb | Dp2TybOlig1/2KO or Dp3TybDyrk1aKO | One-way ANOVA F and P value | P Value (WT vs Dp) | P Value (Dp vs KO) | P Value (WT vs KO) |
|---|---|---|---|---|---|---|---|---|
| Dp2TybOlig1/2KO | Frequency (Hz) | 3.6 ± 0.4 (33) | 3.6 ± 0.4 (35) | 3.4 ± 0.4 (26) | 0.03 KW, 0.9825 | ns | ns | ns |
| | Amplitude (pA) | -30 ± 1.3 (33) | -28.5 ± 1 (35) | -30.7 ± 1.2 (27) | 2.1 KW, 0.315 | ns | ns | ns |
| | Tonic current (pA) | 7.9 ± 0.9 (32) | 12.7 ± 1.5 (30) | 8.3 ± 1.3 (27) | 9.5 KW, 0.0084 | P<0.05 | P<0.05 | ns |
| | Capacitance (pA) | 89.8 ± 3.7 (33) | 89.8 ± 4 (35) | 82.5 ± 4.5 (27) | 0.98, 0.3799 | ns | ns | ns |
| Dp3TybDyrk1aKO | Frequency (Hz) | 2.9 ± 0.3 (25) | 3.2 ± 0.4 (38) | 2 ± 0.2 (38) | 4.2 KW; 0.1197 | ns | ns | ns |
| | Amplitude (pA) | -29.1 ± 1.4 (25) | -29.5 ± 1 (38) | -29 ± 0.9 (38) | 0.15 KW; 0.9283 | ns | ns | ns |
| | Tonic current (pA) | 9.1 ± 1.1 (23) | 14.6 ± 1.8 (34) | 9.2 ± 1.2 (34) | 4.61, 0.0125 | P<0.05 | P<0.05 | ns |
| | Capacitance (pA) | 86.4 ± 3 (25) | 85.7 ± 4.3 (38) | 90.1 ± 4.1 (38) | 1.1 KW; 0.5781 | ns | ns | ns |

Frequency and amplitude of phasic inhibition along with tonic currents and whole-cell capacitance of duplication (Dp) mouse models compared to littermate wild-type controls. Number in brackets show n numbers of cells. P and F values were derived from one-way ANOVA or Kruskal-Wallis nonparametric ANOVA and can be found in figures and their legends. KO, knockout; KW, Kruskal-Wallis Test; ns, not significant
